# Purine-rich RNA sequences in the 5’UTR site-specifically regulate eIF4A1-unwinding through eIF4A1-multimerisation to facilitate translation

**DOI:** 10.1101/2022.08.08.503179

**Authors:** Tobias Schmidt, Adrianna Dabrowska, Joseph A. Waldron, Kelly Hodge, Grigorios Koulouras, Mads Gabrielsen, June Munro, David C. Tack, Gemma Harris, Ewan McGhee, David Scott, Leo M. Carlin, Danny Huang, John Le Quesne, Sara Zanivan, Ania Wilczynska, Martin Bushell

## Abstract

Oncogenic translational programmes underpin cancer development and are often driven by dysregulation of oncogenic signalling pathways that converge on the eukaryotic translation initiation (eIF) 4F complex. Altered eIF4F activity promotes translation of oncogene mRNAs that typically contain highly structured 5’UTRs rendering their translation strongly dependent on RNA unwinding by the DEAD-box helicase eIF4A1 subunit of the eIF4F complex. While eIF4A1-dependent mRNAs have been widely investigated, it is still unclear how highly structured mRNAs recruit and activate eIF4A1 unwinding specifically to facilitate their preferential translation.

Here, we show that RNA sequence motifs regulate eIF4A1 unwinding activity in cells. Our data demonstrate that eIF4A1-dependent mRNAs contain AG-rich motifs within their 5’UTR which recruit and stimulate eIF4A1 unwinding of localised RNA structure to facilitate mRNA translation. This mode of eIF4A1 regulation is used by mRNAs encoding components of mTORC-signalling and cell cycle progression and renders these mRNAs particularly sensitive to eIF4A1-inhibition. Mechanistically, we show that binding of eIF4A1 to AG-rich sequences leads to multimerization of eIF4A1 with eIF4A1 subunits performing distinct enzymatic activities. Our structural data suggest that RNA-binding of multimeric eIF4A1 induces conformational changes in the RNA substrate resulting in an optimal positioning of eIF4A1 proximal to the RNA duplex region that supports efficient unwinding.

Hence, we conclude a model in which mRNAs utilise AG-rich sequences to specifically recruit eIF4A1, enabling assembly of the helicase-active multimeric eIF4A1 complex, and positioning these complexes proximal to stable localised RNA structure allowing ribosomal subunit scanning.

## INTRODUCTION

Dysregulation of cellular translation is a prominent feature of many cancers supporting proliferative gene signatures and establishing oncogenic programmes initiated through signalling pathways including KRAS and mTORC (Meyer and Penn, 2008; Saxton and Sabatini, 2017). Downstream of these pathways operates a key factor of eukaryotic translation initiation (eIF), the eIF4F complex, the activity of which links oncogenic signalling to oncogenic protein synthesis (Lazaris-Karatzas et al., 1990; Modelska et al., 2015; Wolfe et al., 2014). eIF4F consists of the cap-binding protein eIF4E, the scaffold protein eIF4G and the ATP-dependent DEAD-box RNA helicase eIF4A1 that displays ATPase-dependent RNA strand separation activity. By virtue of the eIF4F-complex, eIF4A1 catalyses at least two major steps in translation: mRNA loading onto the 43S PIC and its translocation along the 5’ UTR to the translation start site (Kumar et al., 2016; Shirokikh et al., 2019; Sokabe and Fraser, 2017; Svitkin et al., 2001; Yourik et al., 2017). Interestingly, the loading function requires only eIF4A1’s ATPase activity (Sokabe and Fraser, 2017; Yourik et al., 2017), while unwinding is additionally critical for efficient translation of mRNAs with highly structured 5’UTRs, which are hence considered highly eIF4A1-dependent and include mRNAs of oncogenes such as *MYC* and *BCL2* (Modelska et al., 2015; Rubio et al., 2014; Waldron et al., 2019; Wolfe et al., 2014). A variety of approaches have been aimed at identifying and characterising eIF4A1’s cellular mRNA targets, which have been shown to have longer and more C/GC-rich 5’UTRs, thus containing more RNA secondary structure (Rubio et al., 2014; Steinberger et al., 2020; Waldron et al., 2019; Wolfe et al., 2014). Yet, it is still unresolved whether such highly structured eIF4A1-dependent mRNAs recruit and activate eIF4A1 unwinding specifically. However, selective inhibition of eIF4A using a variety of natural compounds, including silvestrol, hippuristanol, pateamine A and elatol, have all demonstrated anti-tumour activity through downregulation of eIF4A1-dependent genes (Bordeleau et al., 2006; Cencic and Pelletier, 2016; Peters et al., 2018; Wolfe et al., 2014).

eIF4A1 binds ssRNA in an ATP-dependent manner (Hilbert et al., 2009) and ATP-hydrolysis guides the protein through a conformational cycle providing a model for how ATP-turnover and ssRNA-binding are coupled (Andreou and Klostermeier, 2014; Garcia-Garcia et al., 2015; Nielsen et al., 2011; Rogers et al., 2001). However, while it is understood that eIF4A1 unwinds duplex regions within RNAs, eIF4A1 appears to not associate with dsRNA in a detectable manner (Rogers et al., 1999; Wilczynska et al., 2019), hence it still remains unclear how exactly the strand separation step of the duplex region is realised during the ATPase-driven conformational cycle. Moreover, eIF4A1 is a weak helicase by itself but its unwinding efficiency is strongly stimulated in the presence of the cofactors eIF4G, eIF4B and eIF4H. This is achieved by complex formation between eIF4A1 and the cofactor proteins that synergistically modulate eIF4A1’s conformational cycle (Andreou and Klostermeier, 2014; Feoktistova et al., 2013; Garcia-Garcia et al., 2015; Harms et al., 2014; Hilbert et al., 2011; Nielsen et al., 2011; Rogers et al., 2001; Rogers et al., 1999). Each cofactor is believed to operate at a different step of the cycle but since structural information is lacking, it is unclear how multiple cofactors bind and synergise during a single catalytic cycle. Moreover, the exact role of eIF4A1-cofactors in orchestrating eIF4A1’s function in mRNA-loading and unwinding in the translation of specifically eIF4A1-dependent mRNAs is unclear.

Being an essential translation initiation factor, eIF4A1 is considered to bind and load all mRNAs onto ribosomes regardless of RNA sequence and structure (Sokabe and Fraser, 2017; Yourik et al., 2017). However, more recent evidence suggests that the RNA sequence might influence eIF4A1 function: i) The length of an ssRNA substrate affects activities of the yeast eIF4A-eIF4B-eIF4G complex *in vitro* (Andreou et al., 2019), ii) members of the eIF4A protein-family preferentially bind to distinct mRNA sets (Hauer et al., 2015; Shibuya et al., 2004; Wilczynska et al., 2019) and iii) rocaglamide-compounds induce translational repression by clamping eIF4A1 sequence-specifically onto AG-repeats (Iwasaki et al., 2016). Despite this, the role of the RNA substrate itself in regulating eIF4A1 function has not been investigated in detail. We still do not know exactly how different RNA sequences interact with eIF4A1 and its cofactors and how this impacts eIF4A1’s function in translation initiation. Therefore, we set out to investigate the central question whether RNA sequences regulate eIF4A1 activity and function.

Here, we show that eIF4A1 function in cells is regulated by ssRNA sequences in the 5’UTR of mRNAs. We find that eIF4A1 interacts with ssRNA in a sequence-dependent manner involving a process in which eIF4A1 multimerises particularly on AG-rich RNA sequences. Our data shows that eIF4A1-multimerisation stimulates site-directed unwinding of local RNA structure to specifically facilitate translation of otherwise repressed mRNAs. mRNAs that use this mechanism of eIF4A1 regulation encode for components of cell cycle regulation and mTORC-signaling. Our model of eIF4A1 regulation by ssRNA sequences is supported by a) *in vitro* experiments demonstrating eIF4A1 performing RNA sequence-specific activities that are most stimulated by AG-repeat sequences, b) a transcriptome-wide analysis revealing that mRNAs containing AG-repeat motifs in their 5’UTRs show pronounced gain of RNA structure in their 5’UTR and display strongly reduced translation rates following inhibition of eIF4A1 with hippuristanol, and c) a mechanistic investigation showing that eIF4A1 multimerises upon binding to AG-rich ssRNA sequences, directly loading eIF4A1 onto proximal RNA structures and thus activating unwinding. Altogether, our data demonstrate that AG-RNA sequences regulate eIF4A1 function to drive translation of eIF4A1-dependent mRNAs with localised repressive RNA structures, including mRNAs critical for cell cycle progression.

## RESULTS

### eIF4A1 unwinding and stimulation of translation is RNA sequence-specific *in vitro*

eIF4A1 is considered to bind RNA sequence-unspecifically (Henn et al., 2012; Linder and Jankowsky, 2011). However, a more recent study by Iwasaki *et al*., in which short RNAs from a library bound by recombinant eIF4A1 were sequenced after immunoprecipitation (bind-n-seq) (Iwasaki et al., 2016), suggested sequence preferential binding of eIF4A1 (**Supplementary Fig. 1A**), but the effect of the ssRNA sequences on the catalytic capacities of eIF4A1 was not investigated. To examine this, we first measured the RNA unwinding and ATPase activity of recombinant eIF4A1 (**Supplementary Fig. 1B**) *in vitro* using RNA substrates containing an identical 24 bp duplex with 20 nt 5’ overhang sequences that were expected to span a range of RNA binding affinities based on the bind-n-seq experiment by Iwasaki *et al*. (**Fig. 1A**) (Iwasaki et al., 2016). This not only validated RNA sequence specific binding of eIF4A1, but also showed ssRNA sequence-dependent unwinding activity that was most stimulated by AG-repeats and the least stimulated by UC-repeats (**Fig. 1A**). RNA sequence-specific unwinding was also observed in the presence of cofactors eIF4H and eIF4G (**Fig. 1B**), while differential RNA sequence-specific affinities of eIF4A1 were almost abolished. In comparison the ssRNA overhang sequence had nearly no effect on eIF4A1’s ATPase activity (**Supplementary Fig. 1C**), hence, ssRNA sequences mainly influenced unwinding by eIF4A1 which could not entirely be explained by differential RNA binding affinities nor ATPase activities alone (**Fig. 1B**).

**Figure 1.**
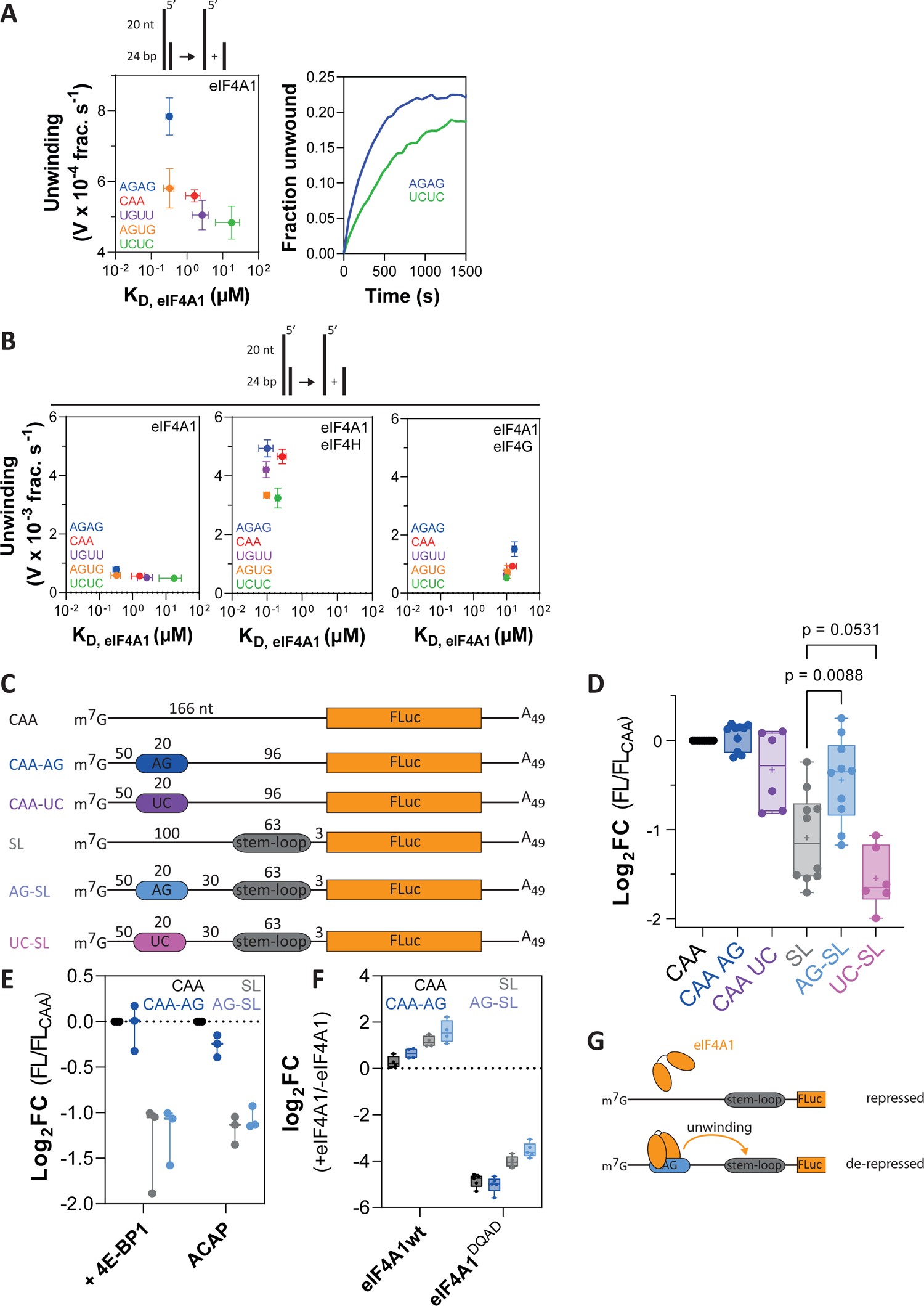
RNA sequence-specific unwinding of eIF4A1 *in vitro*. **A,** ATP-dependent unwinding by eIF4A1 using 5’ overhang-24bp substrates with the indicated 20 nt repeat overhang sequences and RNA-binding of eIF4A1 to the same 20 nt repeat ssRNA in the presence of AMP-PNP (left panel). Data are mean ± SEM from repeat experiments, n = 4. Right panel, example progress curves of eIF4A1 unwinding using the indicated substrates. **B,** same as Fig. 1A (left panel) and unwinding and RNA binding affinity by eIF4A1 in the presence of eIF4H (middle panel) or eIF4G (right panel). Data are mean ± SEM from repeat experiments, n(eIF4H, binding) = n(eIF4H, unwinding) =3, n(eIF4G, binding)=3, n(eIF4G, unwinding)=5. **C,** Firefly luciferase (FLuc, FL) reporter constructs used in *in vitro* translation assays in untreated rabbit reticulocyte lysate (RRL) shown in d-f. **D-F,** relative reporter translation rates measured by the maximum increase in FL-activity over the course of 60 min of translation of indicated reporters in nuclease-untreated RRL at indicated conditions normalised to respective CAA-reporter. Data are mean ± SEM from repeat experiments; **D**, n ≥ 6; **E**, n(+4EBP1) = 3, n(ACAP) = 3. p-values calculated by two-tailed t-test. **F**, log2-fold change of reporter translation rates of indicated reporters in the presence of indicated recombinant eIF4A1 variants. Data are mean ± sem from repeat experiments, n = 4. **G**, schematic presentation of hypothesised AG-motif dependent activity of eIF4A1.

We next asked, if this differential, sequence-dependent unwinding has a functional impact on translation. For this, we employed luciferase-reporter translation assays *in vitro* to specifically examine the effect of the tested sequences with the largest differential in unwinding *i.e.,* AG- and UC repeats. Assays were performed in nuclease-untreated rabbit reticulocyte lysate (RRL) with capped mRNA constructs that contained a linear 5’UTR of CAA-repeats ± a double stem-loop (SL, 2x 11 bp and 4 nt loop) ± a 20 nt AG- or UC-repeat positioned upstream of the SL (**Fig. 1C**). While the SL effectively repressed translation, the presence of an AG-box upstream of the SL, strikingly, led to de-repression (**Fig. 1D**). In contrast, no such effect was observed on the translation of the linear reporters (**Fig. 1D**) nor when an AG-repeat RNA was added in *trans* to the SL-reporter (**Supplementary Fig. 1D**), nor with the UC sequence instead (**Fig. 1D**). Altogether this strongly suggested that de-repression of the structured reporter by the AG-repeat is specific (**Fig. 1D**). This was supported by using RNA substrates corresponding to the reporter 5’UTRs, showing that both eIF4A1 binding affinity and unwinding was stronger if the 5’UTR contained the AG-repeat (**Supplementary Fig. 1E-G**).

To investigate if AG-dependent de-repression is cap-dependent, we added recombinant 4E-BP1, an eIF4E-cap binding inhibitor to the reactions, or used mRNA constructs with a non-functional ApppG-cap (A-cap), which both showed that de-repression by site-specific unwinding requires cap-dependent translation initiation (**Fig. 1E**). To next examine the specific role of eIF4A1 for AG-dependent de-repression, we added recombinant eIF4A1 wild-type (eIF4A1^wt^) or eIF4A1^E183Q^, which is catalytically inactive (hereafter named eIF4A1^DQAD^, **Supplementary Fig. 1H**) (Pause and Sonenberg, 1992) to the reactions. While addition of eIF4A1^wt^ stimulated translation particularly of the SL reporters, eIF4A1^DQAD^ strongly inhibited translation of all reporters demonstrating the strict dependency of reporter translation of eIF4A1 (**Fig. 1F**).

Concluding, eIF4A1 interacts with ssRNA in a sequence-specific manner, which results in RNA-specific activation of RNA unwinding that favours translation of structurally repressed reporter mRNAs in cap-dependent translation *in vitro* (**Fig. 1G**).

### RNA sequence-specific unwinding and simulation of translation by eIF4A1 in cells

To examine the global connection between primary RNA sequence and eIF4A1-dependency of specific mRNA on translation in cells, we applied metabolic pulse-labeling together with quantitative TMT labelling (TMT-pSILAC) in MCF7 cells over a time course immediately following inhibition of eIF4A1 with hippuristanol, which prevents eIF4A1 RNA-binding and unwinding (Bordeleau et al., 2006; Cencic and Pelletier, 2016) (**Fig. 2A**). To measure the associated change in translation per protein in response to eIF4A1-inhibition, we calculated the apparent translation rate of newly synthesized proteins (*k*_hippuristanol_, *k*_DMSO_, **Fig. 2B**). The experiment was performed in quadruplet which uniquely allowed us the capacity to confidently measure direct changes in protein synthesis rates following eIF4A1-inhibition (**Supplementary Fig. 2A-B**). In agreement with hippuristanol being a translational inhibitor (Bordeleau et al., 2006; Cencic and Pelletier, 2016; Waldron et al., 2019), translation rates were nearly exclusively downregulated in response to the treatment (**Fig. 2B**), analysis revealing 255 hippuristanol-sensitive/eIF4A1-dependent mRNAs (254 repressed, 1 upregulated) and 244 hippuristanol-resistant/eIF4A1-independent mRNAs (**Fig. 2C-D, Supplementary Fig. 2C-D**).

**Figure 2.**
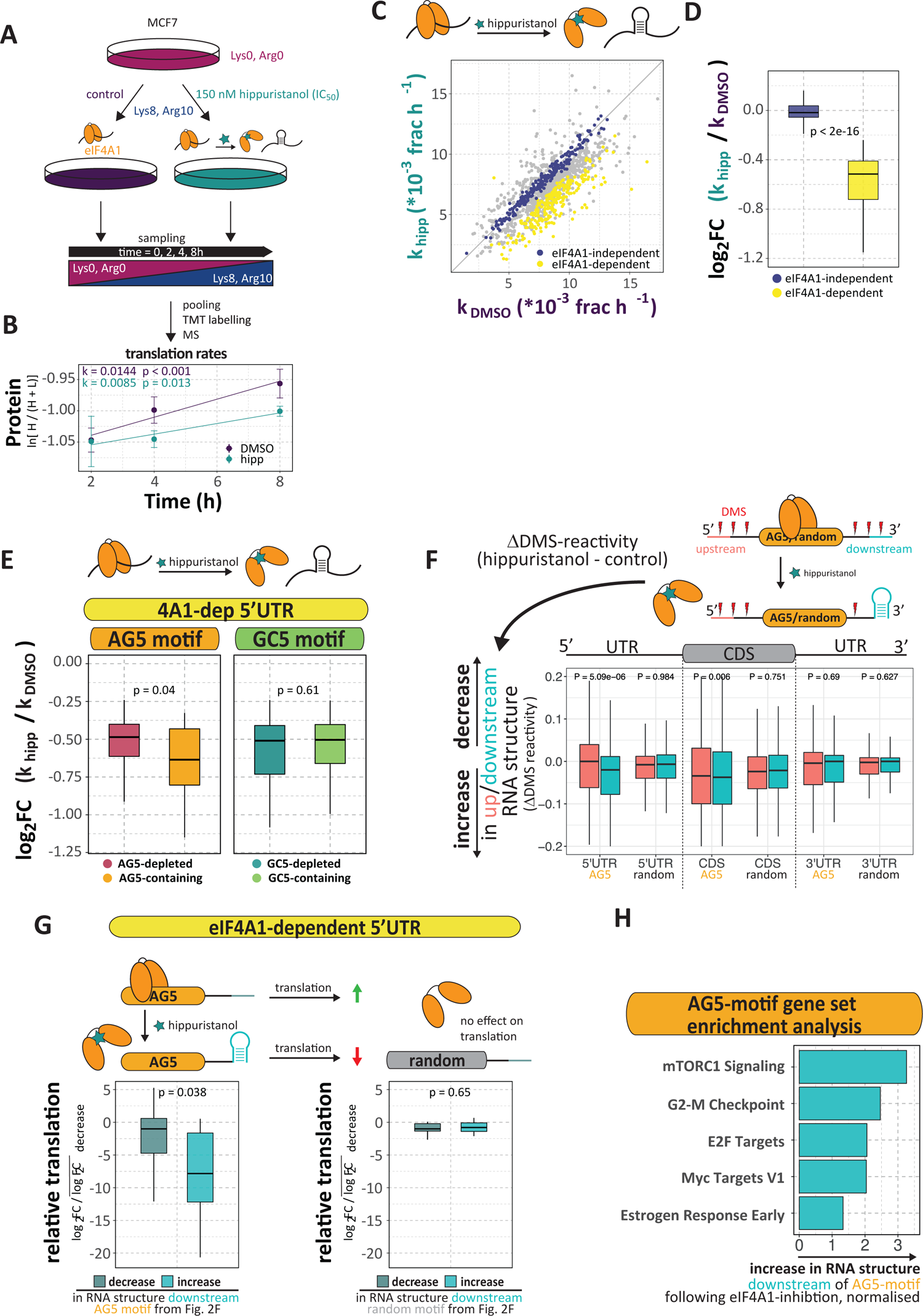
RNA sequence-specific unwinding of eIF4A1 in cells. **A**, Schematic presentation of the performed quantitative TMT pulsed SILAC in MCF7 cells following inhibition of eIF4A1 with hippuristanol. Four independent replicates of labelling experiments were analysed. **B**, Production of newly synthesized KPNB1 over time under control and hippuristanol-treated conditions measured as the natural logarithm of the fraction of incorporation of the (Lys8, Arg10)-labelled (H) protein over total protein. The apparent translation rate *k* of newly synthesized protein is given as the slope of a linear fit. p-values of F-test against k = 0. **C**, Scatter plot of translation rates under control and hippuristanol conditions. eIF4A1-dependent: FDR < 0.1, eIF4A1-independent: FDR > 0.7. **D**, Box plot of the log2-fold change in translation rate following eIF4A1 inhibition of eIF4A1-dependent and – independent mRNAs. **E**, Box plot of the log2-fold change in translation rate following eIF4A1 inhibition of eIF4A1-dependent mRNAs whose 5’UTRs contain AG5- or GC5-motifs. **F**, Box plot of the change in DMS-reactivity (ΔDMS) of 20 nt windows at positions 31-50 nts up- and downstream of all AG5 motifs and randomly selected motifs from 5’UTRs, CDSs and 3’UTRs from the same transcripts. p-values were calculated by a paired, two-sided Wilcoxon test. **G**, Box plots and schematic presentation of the association of the log2-fold change in translation rate and the change of RNA structure in RNA regions 31-50 nt downstream AG5-motifs or random locations of eIF4A1-dependent mRNAs (same RNA regions as Fig. 2F). **H**, Gene set enrichment analysis (hallmarks) of AG5-motif containing mRNAs that show increase of RNA structure downstream of the motif (see Fig. 2F), ranked by increase in RNA structure upon eIF4A1 inhibition (DMS reactivities). FDR of terms < 0.05. Individual group sizes of the panels of this figure are summarized in **Supplementary Table 1.** p-values Fig. 2D-G were calculated by an unpaired or paired, two-sided Wilcoxon test, respectively.

Previous data sets, that investigated the change in translational efficiency following eIF4A1-inactvation, highlighted global 5’UTR features including length, stability and GC-content as markers rendering mRNA translation eIF4A1-dependent (Ho et al., 2021; Iwasaki et al., 2016; Steinberger et al., 2020; Waldron et al., 2019; Wilczynska et al., 2019; Wolfe et al., 2014). Interestingly, these features were not different between eIF4A1-depedent and –independent mRNAs (**Supplementary Fig. 2E**). Additionally, examining the AG-content within the 5’UTRs of these mRNA also showed no global distinction between the two groups of mRNAs (**Supplementary Fig. 2E**). Taken together, this suggested that other, less global mRNA features are responsible for eIF4A1-dependence.

To investigate the role of RNA sequence motifs for eIF4A1-dependent translation, we asked specifically if the presence of AG-sequence motifs within the transcript is associated with differential translation rates similar to what we observed in the *in vitro* experiments (Fig. 1D). For this, we grouped mRNAs if their 5’UTRs contained non-overlapping 10 nt AG motifs (see methods) or, as a reference, GC-repeats (GC5), a previously highlighted marker for eIF4A1-dependence of translation (Rubio et al., 2014; Steinberger et al., 2020; Waldron et al., 2019; Wolfe et al., 2014). This showed that translation rates of eIF4A1-dependent mRNAs with AG5-motifs in their 5’UTRs were significantly stronger repressed after eIF4A1-inhibition (**Fig. 2E**), while, in contrast, the reference mRNA group with GC5-motifs within their 5’UTR was not associated with a change in translation rate upon eIF4A1-inhibition. This suggested that presence of AG5-*motifs* in the 5’UTR of eIF4A1-dependent mRNAs increases their requirement of eIF4A1 activity for translation.

We then asked if the stronger translational repression of AG5-motif containing eIF4A1-dependent mRNAs is related to structural rearrangements induced by inhibition of eIF4A1 activity with hippuristanol treatment. For this, we first wanted to understand if the eIF4A1-dependent changes in translation rates are generally associated with changes in RNA structure. To do so, we took advantage of our previous Structure-seq2 data (Waldron et al., 2019) that have also been obtained in MCF7 cells following specific inhibition of eIF4A1 with hippuristanol (**Supplementary Fig. 2F**) (Bordeleau et al., 2006; Cencic and Pelletier, 2016). To evaluate the change in RNA structure, we compared the ΔDMS-reactivity (hippuristanol - control), *i.e.* the change in single-strandedness, of eIF4A1-dependent and –independent mRNAs. This revealed a comparable change in global RNA structure upon eIF4A1-inhibtion between the eIF4A1-dependent and -independent mRNAs within the 5’UTR as well as in the CDS and the 3’UTR (**Supplementary Fig. 2F**). This agrees with our previous findings that eIF4A1-inhibtion does not affect mRNA structure globally (Waldron et al., 2019).

Further, previous studies, including ours examining specifically eIF4A1 (Waldron et al., 2019), have shown that DEAD-box RNA helicases rearrange localised RNA structures (Guenther et al., 2018; Linder and Fuller-Pace, 2013). To specifically test whether AG5-motifs guide local unwinding of RNA structure in an eIF4A1-dependent manner, we compared the change in RNA structure (ΔDMS-reactivity) in 20 nt sliding RNA regions up- and downstream of AG5 motifs (**Supplementary Fig. 2G**). The analysis revealed that in the 5’UTR the content of RNA structure in RNA regions downstream of AG5 motifs increases significantly upon eIF4A1-inhibition (**Fig. 2F** and **Supplementary Fig. 2G**), while this is not observed for RNA regions upstream (**Fig. 2F** and **Supplementary Fig. 2G)** or around randomly selected non-AG5 motifs within the same 5’UTRs (**Fig. 2F**). Neither were site-specific changes in RNA structure observed for RNA regions around AG5-motifs in the CDS or 3’UTR of the same transcripts (**Fig. 2F**). Interestingly, the location of the AG5 motifs in the 5’UTR was unbiased (**Supplementary Fig. 2H**) and the stabilities of RNA structures folded from the DMS-reactivities of the RNA regions downstream of these 5’UTR-AG5 motifs were not different from the stabilities calculated from random locations within the same 5’UTR (same AG5 and random regions as in Fig. 2F, **Supplementary Fig. 2I**). Altogether, this strongly suggested site-specific eIF4A1-dependent unwinding downstream the AG5 motifs in the 5’UTR of eIF4A1-dependent mRNAs (scheme **Fig. 2F**).

Finally, we asked if changes in RNA structure in these eIF4A1-unwinding dependent RNA regions downstream of the AG5 motifs in the 5’UTR affected translation of the mRNA (see also Fig. 2E). For this we paired the change in RNA structure of these RNA regions (Fig. 2F) with the change in translation rate (Fig. 2D) of the respective mRNA following eIF4A1-inhibition. This revealed that gain of RNA structure in RNA regions downstream of the 5’UTR AG5 motifs was associated with translational repression following eIF4A1-inhibition (**Fig. 2G**, same AG5 motifs and regions as in Fig. 2F). In contrast, this was not the case for random locations within the 5’UTR of the same transcript (**Fig. 2G**, same random motifs and regions as in Fig. 2F) nor RNA structure upstream of these motifs **(Supplementary Fig. 2J**). Thus, this strongly suggested that AG5-motifs stimulate eIF4A1-dependent unwinding of downstream RNA structure to facilitate mRNA translation (scheme **Fig. 2G**). Gene set enrichment analysis suggests that mRNAs containing AG-motif that use this mechanism to activate eIF4A1 play a critical role in the translation of known mRNAs with proliferative signature including components of mTORC-signalling and cell cycle progression as well as myc targets (**Fig. 2H**). Taken together, AG-rich RNA sequences in the 5’UTR site-specifically regulate eIF4A1 helicase activity to facilitate translation of eIF4A1-dependent mRNAs with local repressive RNA structure, including mRNAs critical for cell cycle progression.

### RNA sequence-dependent unwinding by eIF4A1 is stimulated by eIF4A1-multimerisation

To assess how RNA sequences, in particular the AG-repeat sequences, specifically activate and stimulate eIF4A1 unwinding mechanistically, we next examined eIF4A1’s catalytic capacities in more detail *in vitro.* For this we aimed to characterise the differential unwinding of substrates with AG and CAA-overhang, to which eIF4A1 displayed comparable affinities (**Fig. 1A**). Titrations confirmed similar functional binding affinity (K_1/2_ ∼ 2 µM, **Fig. 3A**) and that unwinding activity on the CAA-overhang substrate was weaker compared to the AG-overhang (**Fig. 3A**). The functional binding isotherms of the curves for both overhang-sequences were sigmoidal and revealed a Hill-coefficient (*h*) > 1 (**Fig. 3A**). This was also observed at different substrate concentrations and duplex lengths (**Supplementary Fig. 3A-B**) indicating cooperation of multiple eIF4A1 copies in the unwinding reaction, while, in contrast, the ATPase activity did not appear to require cooperation (**Supplementary Fig. 3C**, h ≥ 0.5).

**Figure 3.**
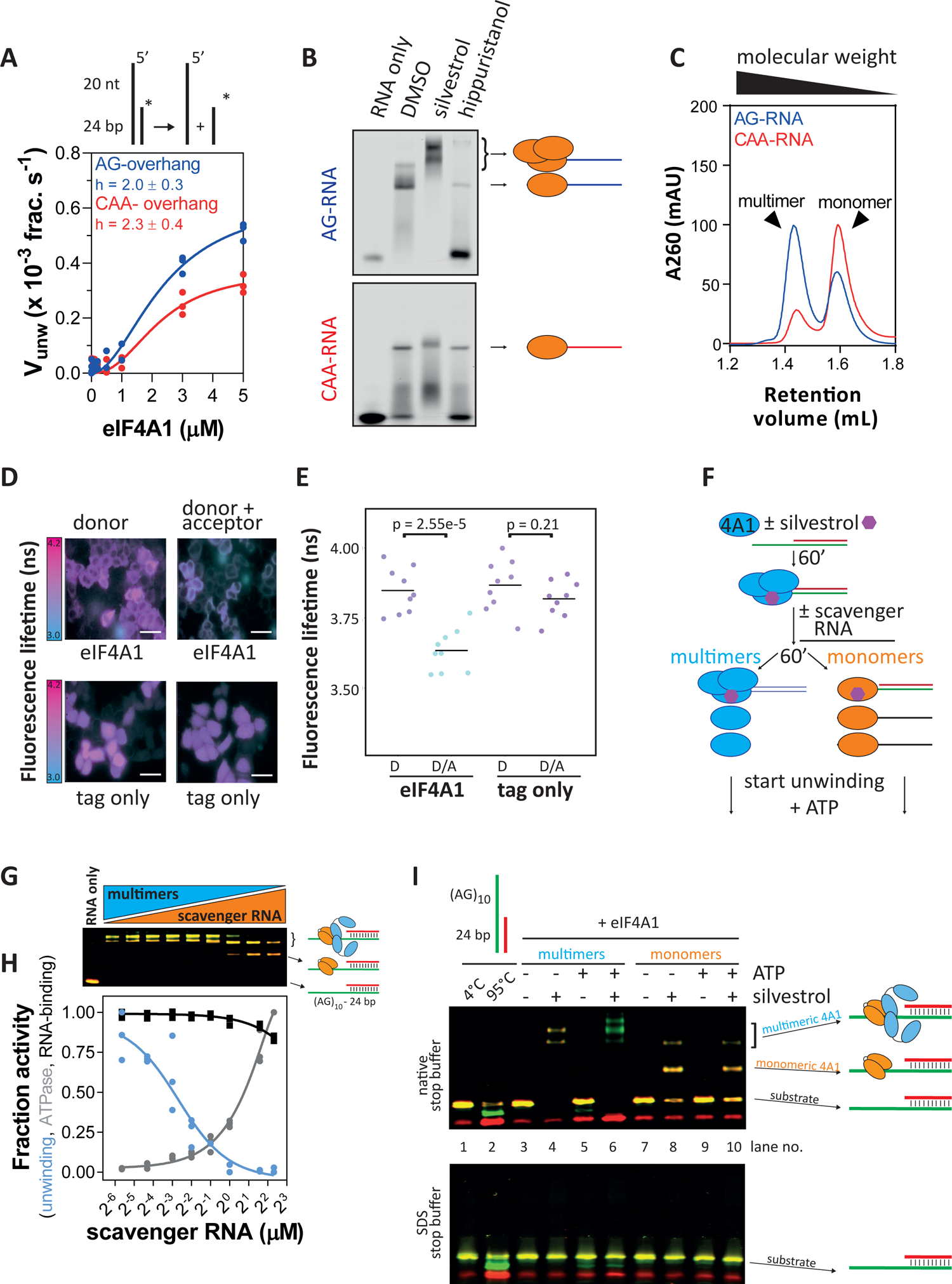
RNA sequence-specific unwinding of is stimulated by RNA sequence-dependent eIF4A1-multimerisation. **A**, Unwinding by eIF4A1 on AG- and CAA-overhang substrates. The Hill-equation was fitted to the data (lines); data are means (technical duplicates) from three repeat experiments, n = 3. **B**, EMSAs of eIF4A1-binding to AG- and CAA-RNA in the presence of 100 µM silvestrol or 50 µM hippuristanol, n = 3 repeat experiments. **C**, Analytical gel filtration of complexes between 16 µM eIF4A1 and 4 µM AG-RNA (blue) or CAA-RNA (red). **D,** Representative images and **E** quantification of fluorescence lifetime imaging of mCitrine-(acceptor, A) and mTurquoise-(donor, D)-labelled eIF4A1 in HeLa cells. Dot plot of n = 9 repeat experiments accounting for total of at least 205 cells per condition. p-values were calculated by a two-sided, unpaired t-test. The scale bar is 50 µm. **F**, Scheme of the clamping reactions for unwinding assay in (G-I). **G,** EMSA of 5 µM eIF4A1 binding to 50 nM AG-overhang substrate after addition of scavenger RNA at increasing concentrations corresponding to Fig. 3H. **H,** Results of fluorescence-based unwinding (blue) and ATPase (grey) assay and RNA-binding (black) using the AG-overhang substrate at increasing concentrations of scavenger RNA. The mean activity data (of technical duplicates) from three replicates is plotted relative to eIF4A1 activity in the absence of scavenger RNA, n = 3. The Hill-equation was fitted to the data (lines). **I,** Representative EMSA unwinding assay (of three repeat experiments, n = 3) of 3 µM eIF4A1 on 50 nM AG-overhang substrate under more monomeric or multimeric eIF4A1 conditions in the presence and absence of silvestrol.

Since eIF4A1 only contains one active site for unwinding, enzymatic cooperativity would mean involvement of multiple eIF4A1 molecules in the reaction. We therefore sought to resolve putative multimeric eIF4A1-RNA complexes by native electrophoretic mobility shift assays. Multimeric eIF4A1 complexes were clearly detectable with the unwinding-activating AG-RNA but not detectable with the less activating CAA-RNA (**Fig. 3B**, **Supplementary Fig. 3D**). The eIF4A-inhibitor silvestrol promoted eIF4A1 multimerisation specifically on the AG-RNA but not CAA-RNA (**Fig. 3B**, **Supplementary Fig. 3E)**, while hippuristanol reduced RNA-binding of eIF4A1 to both RNAs and abrogated multimeric complex formation (**Fig. 3B**). Analytical gel filtration and ultracentrifugation revealed that eIF4A1 multimerisation is i) only induced upon RNA-binding (**Supplementary Fig. 3F-G),** ii) most pronounced at excess eIF4A1 concentrations **(Supplementary Fig. 3F-G)** and iii) RNA sequence-specific (**Fig. 3B-C**). The largest state of multimeric eIF4A1-complexes showed a stoichiometry of eIF4A1:AG-RNA of 3:1 as determined from analytical gel filtration and ultracentrifugation (**Supplementary Fig. 3F-G** and **Supplementary Table 2**).

Intracellular eIF4A1 is highly abundant over typical total mRNA concentrations (eIF4A1: 10 – 20 µM (Galicia-Vazquez et al., 2012), total mRNA < 1 µM, see methods) at an estimated eIF4A1:mRNA ratio of 6-50:1 (Tauber et al., 2020), conditions at which we observe multimerisation *in vitro*. We then asked if eIF4A1 multimerises in cells, for which we employed fluorescence lifetime imaging-fluorescence resonance energy transfer (FLIM-FRET (Mastop et al., 2017)). FLIM-FRET from over-expression of a pair of fluorescently-tagged eIF4A1-wildtype (eIF4A1^wt^) suggested close proximity of the eIF4A1 molecules in living Hela cells (**Fig. 3D-E** and **Supplementary Fig. 3H-I**). Over-expression of tagged-eIF4A1^wt^ together with tagged RNA-binding deficient eIF4A1^DQAD^ (**Supplementary Fig. 3J-K**) (Pause and Sonenberg, 1992) showed similar results thus indicating an eIF4A1-eIF4A1 interaction in cells similar to what we observed *in vitro*.

Having detected the ability of eIF4A1 to multimerise on AG-RNA, we next performed a direct comparison of the unwinding activity of substrate-bound eIF4A1 when levels of multimerisation were either high or low. For this, conditions were required to allow binding of eIF4A1 to the substrate prior to the unwinding reaction. To avoid ATP, which is required for eIF4A1 to bind RNA and initiates unwinding, we used silvestrol because it clamps eIF4A1 onto RNA in an ATP-independent manner (**Fig. 3F** and **Supplementary Fig. 3L**). This also allowed us to setup the reaction in a way so that the total protein as well as the substrate-bound concentrations between the conditions matched. Since protein excess is required for eIF4A1-multimerisation (**Supplementary Fig. 3F**), we first clamped eIF4A1 to the AG-substrate under conditions that allow multimer formation and then, to reduce the degree of multimerisation, added subsequently unlabelled AG-RNA to scavenge excess eIF4A1 from the solution and the multimers (**Fig. 3F-G**). This revealed that highly multimerised eIF4A1 was fully active while lowly multimerised eIF4A1 displayed only residual unwinding activity even though the AG-RNA substrate was fully bound to eIF4A1 (**Fig. 3H**). In contrast, the increase in ATPase activity reflected binding of eIF4A1 to the added scavenger RNA, validating the active, functional state of the protein. Interestingly, increasing amounts of scavenger RNA reduced eIF4A1’s unwinding activity already by over 70% before multimerisation of substrate-bound eIF4A1 was reduced, indicating that not only substrate-bound but also un-bound, free eIF4A1 molecules participate in the unwinding reaction (**Fig. 3G-H**). To visualise substrate binding and unwinding simultaneously, we performed a dual-colour gel shift unwinding assay under similar conditions. This confirmed that, under clamping conditions (silvestrol) and in the absence of ATP, eIF4A1 was fully bound to the AG-substrate without unwinding it under both high and low multimerisation conditions (**Fig. 3I lanes 4 and 8**). Yet, only under high multimerisation conditions did addition of ATP induce strand separation (**Fig. 3I, lanes 5+6 *vs* 9+10**) and, in addition, overhang-clamped eIF4A1 strongly stimulated unwinding and remained bound to the overhang strand after the reaction (**Fig. 3I, lane 6 and Supplementary Fig. 3M**).

In conclusion, these data suggest that RNA sequence-specific unwinding by eIF4A1, particularly on AG-RNA, is mediated by the overhang sequence of the substrate, allowing eIF4A1-multimerisation that enables cooperation between overhang-bound and -unbound eIF4A1 molecules. These effects are enhanced by AG-repeat sequences.

### Different subunits within the eIF4A1-multimer operate distinctly to enable RNA sequence-specific unwinding

To understand the functional connection between overhang-bound and unwinding-performing subunits within the multimeric eIF4A1 complex better, we aimed to probe for functional cooperativity within the eIF4A1 multimer directly. For this, we followed an approach that has been used for multimeric ATPases previously (Moreau et al., 2007; Werbeck et al., 2008), in which, briefly, the catalytic activity of the multimeric enzyme is monitored when wildtype and an inactive variant are mixed at different fractions but at the same total protein concentration. Absence of functional cooperativity would result in a linear trend, with x + y = 1 (Moreau et al., 2007), plotting activity *versus* fraction of the inactive variant. Mixing eIF4A1^wt^ with catalytically inactive eIF4A1^DQAD^ (**Supplementary Figs. 1H and 3K)** (Pause and Sonenberg, 1992), our results show a differential level of functional cooperativity for eIF4A1 unwinding between the AG- and CAA-overhang (**Fig. 4A**), which correlates with differential unwinding activity on these substrates. This suggested specific activation of eIF4A1 unwinding underlies enhanced functional cooperativity between eIF4A1-subunits within the multimeric eIF4A1 complex in an overhang sequence-dependent manner.

**Figure 4.**
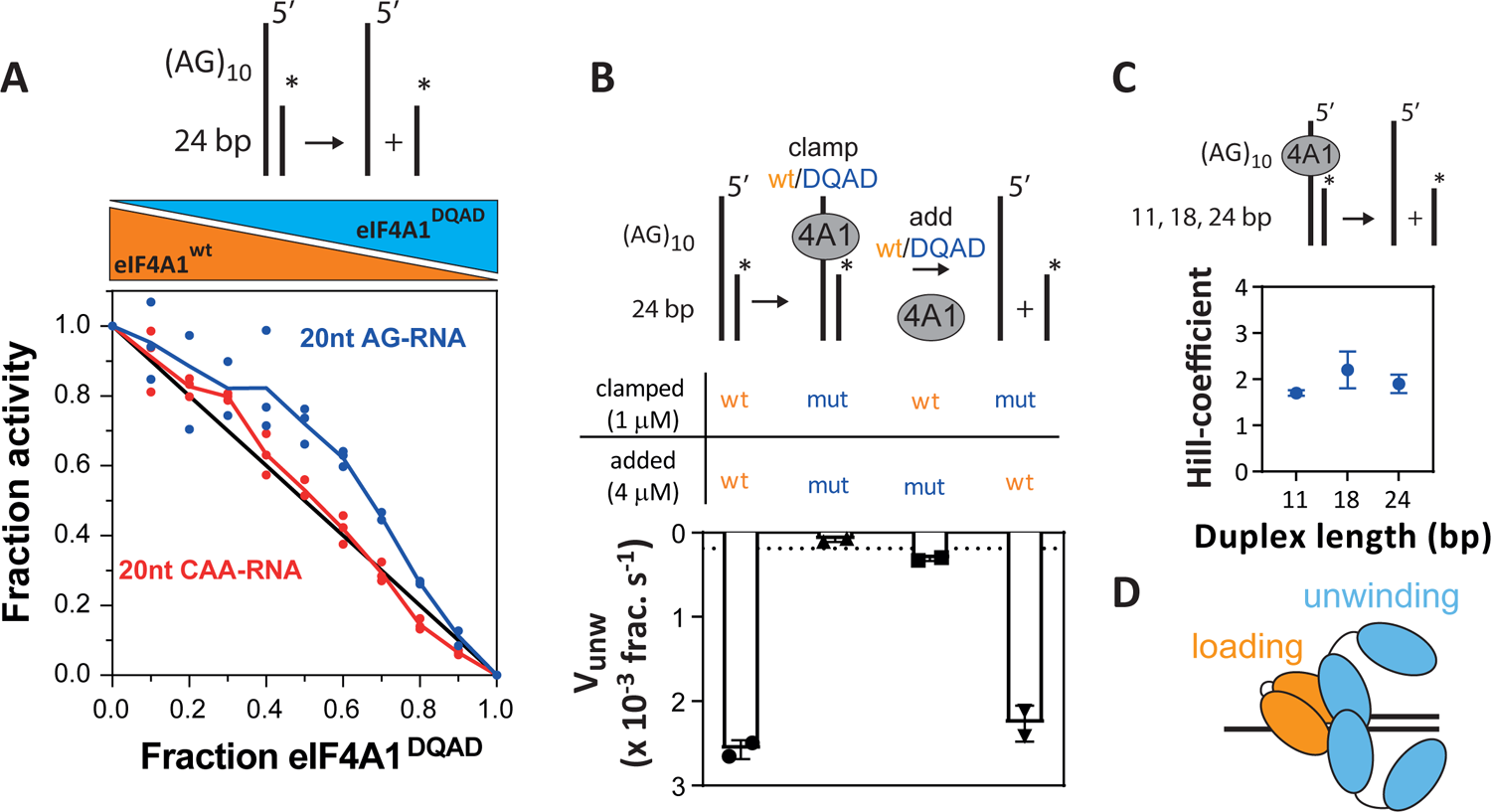
Different subunits within multimeric eIF4A1 perform different functions in RNA-binding, ATPase and unwinding. **A**, Inhibition of unwinding activity on AG-(blue) and CAA-overhang substrate (red) of eIF4A1^wt^ by fractional mixes with inactive variant eIF4A1^DQAD^. The mean activity data (of technical duplicates) from three replicates is plotted relative to non-inhibited eIF4A1 (eIF4A1^wt^ only); data are mean ± sem, n = 3 repeat experiments. The line (x + y = 1) indicates the behaviour of a monomeric or multimeric enzyme without subunit cooperativity. Dashed lines indicate the trend. **B,** Unwinding activities of 5 µM mixed eIF4A1^wt^-eIF4A1^DQAD^ complexes; data are mean ± SD from repeat experiments, n = 2. The dashed line indicates unwinding activity of 5 µM eIF4A1^wt^ on blunt-end substrate. **C,** Hill coefficients of unwinding by clamped eIF4A1^wt^ on different duplex lengths. Data points and errors are results from fitting the Hill equation to the experimental data. **D,** Model of unwinding-competent multimeric eIF4A1.

We next asked if functional cooperativity between eIF4A1 subunits stems from participation of the overhang-bound eIF4A1 directly in the strand separation reaction. For this we asked if binding of catalytically inactive eIF4A1^DQAD^ to the overhang of the substrate before addition of eIF4A1^wt^ inhibits or activates the helicase activity of eIF4A1^wt^ (**Fig. 4B**). To do this, we clamped eIF4A1^DQAD^ first to the overhang of the substrate using silvestrol, which recovered wildtype-like RNA-binding affinity as well as kinetic stability without rescuing its unwinding activity (**Supplementary Figs. 4A-C**). Thus, this excluded the possibility that additional eIF4A1^wt^ could replace overhang-clamped eIF4A1^DQAD^ during the experiment. Strikingly, clamping inactive eIF4A1^DQAD^ to the overhang supported unwinding by eIF4A1^wt^ at a rate similar to the eIF4A1^wt^-only reaction (**Fig. 4B** and **Supplementary Fig. 4D**) indicating beneficial cooperation between the overhang-bound, catalytically inactive eIF4A1^DQAD^ and unwinding-active eIF4A1^wt^. This strongly suggested that the different eIF4A1-copies within multimeric eIF4A1 have different functions. Supporting this, the overall ATPase activity was reduced when eIF4A1^DQAD^ is clamped to the overhang (**Supplementary Fig. 4E**), while unwinding was unaffected (**Fig. 4B**) In this setup, the observed ATPase activity is exclusively performed by eIF4A1^wt^ that performs the actual strand separation, thus, in the wt-only multimeric eIF4A1 complex, the subunits bound to the overhang and subunits performing the strand-separation have different ATPase activities.

In summary, overhang-bound eIF4A1 is not directly involved in unwinding but critical for loading and activating proximal strand separation by distinct eIF4A1 molecules. We thus refer to roles of these different subunits within the multimeric eIF4A1 complex as loading (ssRNA overhang-bound) and unwinding. As Hill-coefficients under clamping conditions indicate activity of two unwinding subunits on short and long duplexes (**Fig. 4C**), we suggest a model in which the eIF4A1-RNA-loading complex activates at least two unwinding subunits (**Fig. 4D**). The catalytic capacity of the loading-complex appears dispensable suggesting a binding-induced mechanism of activation.

### eIF4A1 cofactors operate distinctly upon multimeric eIF4A1

Cellular eIF4A1 function is believed to be tightly regulated through interactions with its cofactors eIF4H, eIF4B and eIF4G (Andreou and Klostermeier, 2014; Garcia-Garcia et al., 2015; Nielsen et al., 2011; Rogers et al., 2001). As our initial results showed that the pattern of RNA sequence-specific unwinding activity of eIF4A1 is differently affected by different eIF4A1 cofactors, we therefore investigated if the cofactors operate upon multimeric eIF4A1. Our **Supplementary Results** demonstrate i) that stimulation of RNA sequence-specific unwinding of eIF4A1 by eIF4G or eIF4H is optimal under conditions that allow multimeric eIF4A1 complex formation, ii) that stimulation by eIF4G or eIF4H occurs in an RNA sequence-specific manner. and iii) that eIF4G and eIF4H operate differently on multimeric eIF4A1, with eIF4G functioning upon or replacing the loading subunit while eIF4H improves activity of the unwinding subunits (for detailed presentation see **Supplementary Results and Supplementary Fig. 5**). Altogether, these results demonstrate that activity of eIF4A1 cofactors differentially stimulates multimeric eIF4A1 complexes to facilitate distinct RNA sequence-specific unwinding activities.

### RNA sequence-specific eIF4A1 complexes

To investigate how the RNA sequence facilitates activation of eIF4A1 unwinding at a structural level, we next examined the shape of eIF4A1-ssRNA complexes using small-angle x-ray scattering (SAXS). Envelope models revealed that apo-eIF4A1 fitted better to eIF4A in an open but not in a closed conformation, which was in agreement with an extended conformation of the apo-protein (**Supplementary Fig. 6A**). Moreover, the shape of the eIF4A1-CAA-RNA complex (eIF4A1 bound to CAA-RNA) suggested a similarly extended conformation as observed with apo-eIF4A1, while the eIF4A1-AG-RNA complex (eIF4A1 bound to AG-RNA) was in a different, more compact conformation as compared to eIF4A1-CAA-RNA and apo-eIF4A1 (**Fig. 5A** and **Supplementary Fig. 6B-C** and **Supplementary Table 3**). In support, linear free energy relationship measurements (Schmidt et al., 2016) demonstrated a higher proportion of both ionic and non-ionic interactions in the eIF4A1-AG-RNA complex than in the eIF4A1-CAA-RNA complex suggesting distinct and RNA specific eIF4A1-AG- and eIF4A1-CAA-RNA binding interfaces (**Supplementary Fig. 6D**). Since it has been shown recently that RNA length modulates the conformation of yeast eIF4A (Andreou et al., 2019), this altogether strongly suggested that interactions of human eIF4A1 with RNA length and sequence guide specific conformational transitions in the protein.

**Figure 5.**
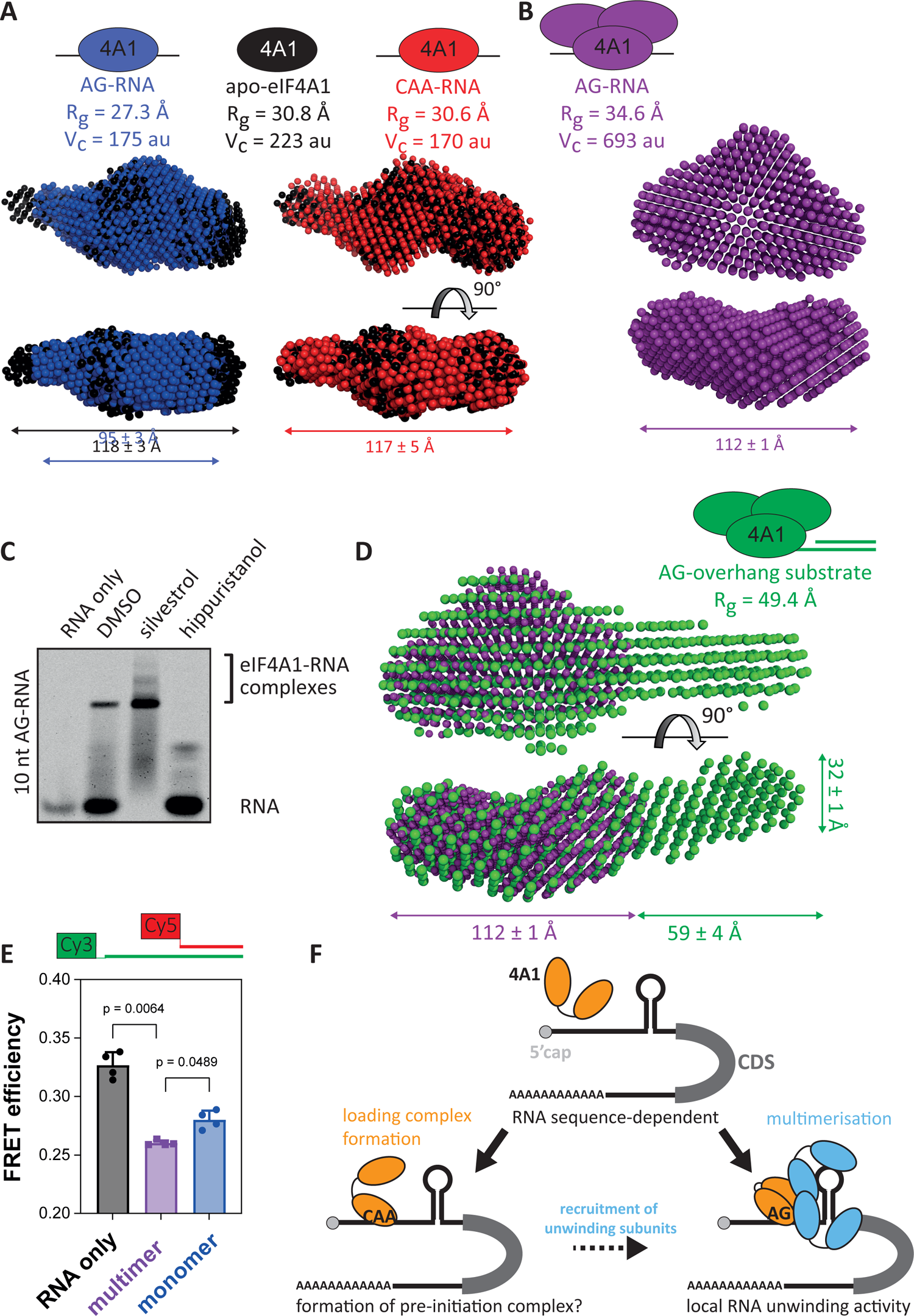
RNA sequence-specific eIF4A1-RNA complexes. **A, B, D**, Small-angle x-ray scattering-based (SAXS) envelope model of **A** apo-eIF4A1 (ligand-free, black), monomeric eIF4A1 bound to AG-(blue) or CAA-RNA (red), respectively, **B** multimeric eIF4A1 bound to AG-RNA (purple) or **D** AG-overhang substrate (green), respectively. Monomer and multimers were generated according to results shown in Fig. 3. V_C_ and R_g_ – Envelope volume of correlation and radius of gyration derived from experimental data; D_max_ – maximum distance is mean ± sd from four measurements based SAXS models using PyMOL2. P(r)-derived Dmax is summarised in Supplementary Table 3. **C,** Representative EMSA of 3 µM eIF4A1 and 10 nt labelled-AG-RNA in the presence of eIF4A1-inhibitors, n = 3 repeat experiments. **E,** Changes in FRET efficiency of labelled AG-overhang substrate upon multimeric and monomer eIF4A1-binding using 0 and 6 µM competitor RNA, respectively (see Fig. 3G); data are mean ± SD from 4 repeat experiments, n = 4. p-values were calculated by one-way ANOVA for paired data. **F,** Model of RNA sequence-regulated activities of eIF4A1. eIF4A1 adopts RNA sequence-specific conformations upon RNA-binding allowing either loading complex formation or multimeric complex formation. mRNAs with AG-rich sequences specifically recruit eIF4A1, enabling assembly of the helicase-active multimeric eIF4A1 complex, and positioning these complexes proximal to stable localised RNA structure allowing ribosomal subunit scanning.

SAXS of multimeric eIF4A1-AG-RNA complexes revealed a non-linear shape of the complex with a larger radius of gyration (*R_g_*) and volume of correlation (V_C_) than monomers (**Fig. 5B**, **Supplementary Fig. 6E-G** and **Supplementary Table 3**). Considering an eIF4A1:RNA stoichiometry of greater than one within the eIF4A1-multimers (**Supplementary Fig. 3F-G** and **Supplementary Table 2**), more than one eIF4A1 subunit within multimeric eIF4A1 could be RNA associated. It is unlikely that more than one subunit binds tightly to the ssRNA because: a 10 nt AG-RNA provides only one direct eIF4A1 binding site, as shown by a recent crystal structure of eIF4A1 in complex with the 10 nt AG-RNA and a silvestrol derivate (Iwasaki et al., 2019), but eIF4A1 multimerisation is still observed on a 10 nt AG-RNA in the presence of silvestrol (**Fig. 5C and Supplementary 3G**). In conclusion, RNA sequence is critical to establish specific binding interfaces and thus conformational states of eIF4A1 that allow formation of multimeric complexes in which only one eIF4A1 protein is in tight contact with the ssRNA sequence.

To investigate how eIF4A1-loading complexes activate unwinding, we next performed SAXS on multimeric eIF4A1 complexes bound to the AG-overhang substrate in the presence of AMP-PNP which reflects the loaded, pre-unwinding state (**Fig. 3I lane 4**). Superposition with the multimeric eIF4A1-AG-RNA complex enabled identification of the overhang (eIF4A1 covered) and the duplex region of the substrate (**Fig. 5D** and **Supplementary Fig. 6H-I**). The measured duplex diameter was slightly larger (24 Å *vs* 32 Å ∼ 33%) than expected, indicating an underestimation of dimensions in the envelopes. Surprisingly, the length of the detected duplex region was shorter than the expected length (59 Å/21 bp *vs* 67 Å/24 bp), suggesting that the eIF4A1-loading complex is located precisely at the overhang-duplex fork and may be covering parts of the duplex region. Fitting a 24 bp dsRNA into the envelope suggests ∼ 5 bps might be buried inside the multimeric eIF4A1-loading complex (**Supplementary Fig. 6J**). Moreover, FRET experiments focusing on the overhang-fork region of the substrate demonstrated a conformational change in the RNA upon eIF4A1 loading complex formation specific to the multimeric state (**Fig. 5E and Supplementary 6K-L**).

Taken together our data support a mechanism in which eIF4A1 undergoes RNA sequence-specific conformational changes that trigger assembly of multimeric eIF4A1-RNA complexes. Within the multimers, the RNA overhang region adopts a conformation that places eIF4A1 subunits directly at the overhang-fork region and partially onto the duplex region. We hypothesise that this is critical for activation of sequence-specific unwinding.

## DISCUSSION

The DEAD-box RNA helicase eIF4A1 catalyses at least two major reactions in translation initiation. First, eIF4A1 activity is essential to load mRNAs onto the 43S pre-initiation complex (PIC) and, second, eIF4A1-dependent unwinding of RNA secondary structure facilitates translocation of the PIC along the mRNAs’ 5’ UTR with high structural content (Kumar et al., 2016; Shirokikh et al., 2019; Sokabe and Fraser, 2017; Svitkin et al., 2001; Yourik et al., 2017). In this study we uncover a mechanism for how such eIF4A1-dependent mRNAs specifically recruit and activate eIF4A1 unwinding activity. Our data reveal that, *in vitro* and in cells, i) eIF4A1 helicase activity is induced in an RNA sequence-specific manner through eIF4A1-multimerisation and ii) that this mechanism of eIF4A1 regulation is used by eIF4A1-dependent mRNAs to overcome translational repression due to localised RNA structure (**Fig. 5F**). Within the 5’UTR of eIF4A1-dependent mRNAs, we identify specific RNA sequence motifs, particularly enriched for polypurines, which function to specifically recruit and trigger eIF4A1-multimerisation to activate eIF4A1-dependent strand separation of local repressive RNA structure to facilitate mRNA translation.

In order to examine the relationship between eIF4A1-dependent unwinding and translation in cells, we combined RNA-structure-seq2 with TMT-pulsed SILAC following eIF4A1 inhibition. In contrast to previous studies, which typically consider results of single time points representing pre- or steady states limiting dynamics, (Steinberger et al., 2020; Waldron et al., 2019; Wolfe et al., 2014) we performed a time course experiment to directly quantify translation rates of newly synthesised proteins immediately after eIF4A1-inhibition (**Fig. 2A-B**). This identified hippuristanol-sensitive/eIF4A1-dependent and hippuristanol-resistant/eIF4A1-independent mRNAs (**Fig. 2C**). Our study finds that eIF4A1-dependent mRNAs do not have longer-than-average 5’UTRs nor increased GC content in contrast to previous reports (Steinberger et al., 2020; Waldron et al., 2019; Wolfe et al., 2014). Proteins identified in our study are detected by a threshold minimum rate of incorporation of the metabolic labelling agent, hence mRNA groups identified through the analysis exhibit fast translation rates naturally. This allowed us to identify features of eIF4A1-dependent mRNAs that increase their sensitivity to eIF4A1-activity. This was achieved through analyses of two independent approaches (RNA structure-seq2 and TMT pulsed SILAC) which revealed RNA sequence-dependent activities of eIF4A1 in cells. This allowed us to define eIF4A1-dependent mRNAs that contain AG-rich motifs in their 5’UTR that essentially facilitate eIF4A1-dependent translation. These findings agree with our *in vitro* data showing that eIF4A1-unwinding activity is stimulated in an RNA sequence-dependent manner, with polypurine-rich sequences enhancing eIF4A1 unwinding the most. mRNAs that contain such AG5-motifs to regulate their translation include well-described eIF4A1-dependent mRNAs such as myc targets, and mRNAs encoding components of cell cycle regulation and mTORC-signalling (**Fig. 2H**). Together, this describes a model in which eIF4A1-dependent mRNAs use AG-rich motifs in their 5’UTR to recruit and specifically activate eIF4A1-unwinding to regulate their translation (**Fig. 5F**).

Mechanistically, our data shows the specific sequence information within the RNA enhances unwinding by eIF4A1 by promoting eIF4A1-multimerisation through an RNA-centric mechanism. Following RNA-sequence specific binding (**Fig. 1A**), eIF4A1 forms ATPase-active but unwinding-inefficient monomeric complexes (**Fig. 3G-H**) or unwinding-activated multimeric complexes (**Fig. 3B** and **Fig. 3I**) directed by RNA sequence (**Fig. 5F**). Activation of unwinding is achieved by a specific division of catalytic capacities between the different eIF4A1-subunits, overhang-bound and unwinding subunits, within multimeric eIF4A1 (**Fig. 4B** and **Supplementary Fig. 4E),** such that overhang-bound eIF4A1 does not directly participate in the unwinding step but stimulates duplex separation by additional eIF4A1-subunits (**Fig. 4B**). eIF4A1-multimerisation itself is initiated by a single eIF4A1 binding to the single-stranded overhang of the substrate (**Fig. 5C**) and subsequently undergoing conformational changes that allow recruitment of additional eIF4A1 subunits and thus formation of the multimeric eIF4A1-loading complex (**Fig. 5A-B**). Assembly of this complex changes the conformation of the single-stranded RNA region (**Fig. 5E**) such that eIF4A1 subunits are positioned at the fork of the proximal RNA duplex (**Fig. 5D** and **Supplementary Fig. 6J**). This then allows enhanced engagement of eIF4A1-unwinding subunits with the duplex stimulating strand separation. We hypothesise that eIF4A1-loading subunits transition dynamically into unwinding subunits which enables recruitment of new loading subunits, free eIF4A1, fuelling the unwinding reaction. This would be in agreement with a requirement of free eIF4A1 for efficient unwinding (**Fig. 3G-I**). A change in the conformation of the ssRNA region upon helicase binding and a similar multimerisation model has been described for of the Ded1p/DDX3 family previously (Kim and Myong, 2016; Putnam et al., 2015). However, while Ded1p/DDX3 binds ssRNA regardless of ATP (Iost et al., 1999), eIF4A1’s ssRNA-binding is dependent on simultaneous ATP-binding and thus eIF4A1’s ATPase activity (Pause and Sonenberg, 1992; Rogers et al., 1999). Interestingly, our results show that ATP-turnover of the loading subunits *per se* is not essential for subsequent unwinding (**Fig. 4B** and **Supplementary Fig. 4E**). Together, this suggests that the ATPase activity of the eIF4A1-loading complexes controls their kinetic stability and thus activation of unwinding as opposed to a direct contribution of the ATPase activity to the strand separation reaction itself. This could in part explain the different unwinding activities of eIF4A1 on different overhang sequences. In agreement, silvestrol-clamped eIF4A1 showed increased unwinding activity.

Multimerisation of DEAD-box helicases as a requirement for efficient RNA strand separation has also been reported for the yeast Ded1p, and its human homolog DDX3X, cold-shock activated helicase CshA and heat-resistant RNA-dependent ATPase Hera (Huen et al., 2017; Putnam et al., 2015; Rudolph et al., 2006; Sharma et al., 2017). These studies present a range of modes how helicases multimerise: While DDX3X/Ded1p forms multimeric complexes readily in the absence of RNA (Putnam et al., 2015), complex formation of CshA and Hera is mediated by unique dimerization domains (Huen et al., 2017; Rudolph et al., 2006). Our data shows, that eIF4A1, in contrast, follows a distinct mechanism. eIF4A1-multimer formation is dependent on RNA-binding (**Fig. 3B** and **Supplementary Fig. 3F-G**) and occurs in an RNA sequence-specific manner with polypurine repeats triggering efficient multimerisation (**Fig. 1A, 3B-C**). Importantly, we observe significant eIF4A1-multimerisation *in vitro* at protein concentrations lower than 5 µM which is lower than the cellular eIF4A1 concentration of 10 – 20 µM (Galicia-Vazquez et al., 2012) and thus would strongly support multimerisation of eIF4A1 occurring in cells. In agreement, eIF4A1-multimers are active in cells, we i) visualised direct eIF4A1-eIF4A1 interactions (**Fig. 3D-E**) and ii) reveal RNA sequence-specific unwinding (**Fig. 2F-G**) as well as stimulation of translation by eIF4A1 in cells (**Figs. 2D and 2H**).

In the cellular environment, the majority of eIF4A1 functions is believed to rely on interactions between eIF4A1 and its cofactors including eIF4G, eIF4B and eIF4H, that collectively stimulate eIF4A1’s catalytic capacities (Andreou and Klostermeier, 2014; Garcia-Garcia et al., 2015; Nielsen et al., 2011; Rogers et al., 2001). It has been shown mechanistically that the different cofactors affect the rates of conformational transitions within eIF4A1 thus guiding eIF4A1 through its catalytic cycle. Our results extend the existing models and shows that cofactors also operate efficiently upon eIF4A1-multimers (**Supplementary Fig. 5B-D**). Our data are consistent with a model in which eIF4H and eIF4G operate on distinct eIF4A1-subunits to deliver their function which allows synergistic activation of multimeric eIF4A1. While eIF4H stabilises the loading complex and stimulates activity of the unwinding subunits, eIF4G functions on or replaces the loading subunits (**Fig. 5F**). Additionally, we observe that in the presence of eIF4G the communication between eIF4A1-subunits is strongly reduced (**Supplementary Fig. 5K**) suggesting that eIF4G can replace the eIF4A1-loading subunits. A similar observation has been described for the Ded1p-eIF4G interaction (Putnam et al., 2015). As a consequence, RNA-binding specificities are delivered through eIF4G rather than eIF4A1. In support of our model, i) eIF4G contains two eIF4A1-binding sites which each induce different catalytic properties of eIF4A1 upon binding (Korneeva et al., 2001; Marintchev et al., 2009; Nielsen et al., 2011), and ii) silvestrol affected the activity of cofactor-containing eIF4A1-multimers distinctly, *i.e.* silvestrol inhibited activity of eIF4G-containing multimeric complexes while it stimulated eIF4H-containing eIF4A1-complexes (**Supplementary Fig. 5L**). This suggests that the mode of action of silvestrol to stimulate unwinding is to clamp and stabilise the loading eIF4A1 subunits. As a result, silvestrol inhibits multimeric eIF4A1 complexes that do not contain eIF4A1 loading subunits, like the eIF4G-containing ones, by clamping and thus inactivating an eIF4A1-unwinding subunit. This is in agreement with recent reports showing that rocaglamides appear to specifically reduce the unwinding activity of eIF4E-independent (constitutively active) eIF4F variants (Kommaraju et al., 2020).

Our data specifically suggest that multimeric eIF4A1 is critical for site-specific unwinding of RNA structures to facilitate cap-dependent translation regardless of cofactor activity (**Fig. 2**). Moreover, eIF4A1 and other DEAD-box helicases, have recently been shown to be major regulators of RNA condensation which show helicase-mRNA-specific networks and can regulate translation (Hondele et al., 2019; Tauber et al., 2020). In their study, eIF4A1 was found to resolve RNA condensates in an unwinding-dependent manner. Our model suggests that eIF4A1 would operate on RNA condensates differentially depending on RNA concentration and RNA sequence composition, resolving preferentially those RNA condensates that allow eIF4A1 multimerisation to occur. However, as eIF4A1-cofactors change the specific activities of multimeric eIF4A1, we hypothesise that, in addition to cofactor activity, a variety of RNA sequences might coordinate eIF4A1 function to drive different translational programmes through recruitment and assembly of distinct multimeric eIF4A1-complexes. This concept might also explain the different 5’UTR features of mRNAs that have been described for eIF4A1-dependent mRNAs. Depending on the approach, the networks between the different multimeric eIF4A1-cofactor complexes might be differentially affected highlighting different but specific groups of eIF4A1-dependent mRNAs. Further, as the active concentration of translation initiation factors including eIF4F is i) tightly controlled in cellular programmes like proliferation and differentiation (Galicia-Vazquez et al., 2014; Mamane et al., 2006; Siddiqui and Sonenberg, 2015), ii) can vary between tissues and iii) is often dramatically affected in many different cancers (Ali et al., 2017; Galicia-Vazquez et al., 2012; Oblinger et al., 2016; Raza et al., 2015), regulation and dysregulation of eIF4A1 multimer formation is likely to have a strong impact on the translational landscape of the cell.

Given the strong evolutionary conservation of the DEAD-box helicase core, it is likely that comparable mechanisms of RNA-based activation of unwinding and hence regulation helicase function are found among the entirety of this protein family (Kim and Myong, 2016; Putnam et al., 2015).

## METHODS

### Cell lines

Hela cells were purchased from ATCC for this study and were already authenticated. In-house authentication using Promega GenePrint 10 was also performed and confirmed Hela identity. All cell lines were tested on a two-weekly basis for mycoplasma. All tests were negative and confirmed the absence of mycoplasma contamination.

### Cell culture and transfection for FLIM experiments

HeLa cells were seeded with cell density of ∼120 000 cells per dish (35 mm sterile MatTek, glass bottom) in DMEM (Gibco) supplemented with 10 % FBS (Gibco) and 2 mM final concentration of L-glutamine (Gibco). Cells were transfected with 1 µg of plasmid, or 1 µg each in case of co-transfections, using GeneJammer (Agilent) at a reagent:plasmid ratio of 3:1. At 48 hours post transfection, medium was exchanged for DMEM (Gibco) supplemented with 10 % FBS (Gibco) and 2 mM final concentration of L-glutamine (Gibco) and cells dishes were taken for FLIM measurements.

### Western blotting

Cells were harvested and lysed in RIPA buffer (50 mM Tris/HCl pH 7.5, 150 mM NaCl, 1 % (v/v) Triton X-100 (Merck), 0.5 % (w/v) sodium deoxycholate (Merck), 0.1 % (v/v) SDS, 5 mM DTT, 0.5 mM PMSF, 5 mM NaF and protease inhibitors (complete EDTA-free, Roche). Lysates were cleared by centrifugation and protein concentration quantified with Bradford. Equal amounts of total protein were loaded onto 4-12% gradient NuPAGE Bis-Tris gels (Invitrogen). Proteins were blotted onto 0.45 µm nitrocellulose membrane using wet transfers. Vinculin is the loading control.

### Antibodies for western blotting

Antibodies were diluted into 1x-TBST supplemented with 5 % (w/v) milk. eIF4A1: ab31217 (Abcam); GFP (which detects mCitrine and mTurquoise, ab13970, Abcam); and vinculin: ab129002 (Abcam)

### Biomass production for generation of recombinant proteins

All proteins were heterologously produced in *E. coli* BL21 (DE3) CodonPlus-RP as N-terminal 6xHis-SUMO-fusion proteins, following procedures as reported in our previous work (Wilczynska et al., 2019). Except for eIF4G, recombinant proteins were produced applying standard protocols for IPTG-induction. Briefly, main cultures were inoculated from overnight pre-cultures. Main cultures were then grown to OD600 = 0.8 - 1 before protein production was induced with a final concentration of 1 mM IPTG. Cells were harvested 4h post induction. For eIF4G, cells were first cultivated at 37 °C to an OD600 = 0.6 – 1 before cells were cooled down to 20 °C and protein production induced with IPTG for 16 h. Cells were harvested by centrifugation and stored at −80 °C.

### Protein and plasmid constructs

For generation of pET-SUMO constructs, cDNAs coding for eIF4A1 (primers TS3/TS4) and eIF4G (674-1600, primers TS9/TS10) and eIF4H (primers TS15/TS16) were generated using standard PCR and subsequently cloned into pET-SUMO vector using the BsaI and NotI restriction sites. eIF4G (674-1600, primers TS9/TS10) was cloned via blunt-end/NotI into linearised pET-SUMO that had been PCR-amplified using primers TS1/TS2 and digested with NotI. eIF4A1^DQAD^ (eIF4A1^E183Q^) was generated by site-directed mutagenesis with the primers TS23/TS24 using the pET-SUMO-eIF4A1 construct. For generation of mTurquoise- and mCitrine-constructs, cDNAs coding for eIF4A1 and eIF4A1^DQAD^ were PCR-amplified using primers TS64/TS65 and cloned into mTurquoise-C1 and mCitrine-C1 vectors (Addgene #54842 and #54587) using HindIII and BamHI restriction sites. Then, mTurquoise- and mCitrine-eIF4A1 were subcloned into pET-SUMO by PCR-amplification using TS4/TS74 using BsaI and NotI restrictions sites. Primers are listed in **Supplementary Table 4**.

### Protein purification

Recombinant proteins were purified following procedures as reported in our previous work (Wilczynska et al., 2019). Cells were resuspended and lysed in buffer A [20 mM Tris/HCl, pH 7.5, 1 M NaCl, 30 mM imidazole and 10 % (v/v) glycerol] supplemented with 1 mM PMSF and complete EDTA-free protease inhibitor cocktail (Roche). After centrifugation at 45 000 x g supernatant was filtered (0.45 µm) and applied to HisTrap (GE Healthcare) affinity chromatography. Bound protein was eluted with a linear imidazole gradient. Pooled fractions were diluted in buffer B [20 mM Tris/HCl, pH 7.5, 10 % (v/v) glycerol, 0.1 mM EDTA, 2 mM DTT] and incubated with SUMO-protease over night at 4°C for cleavage of the SUMO-tag. The protein solutions were further diluted with buffer B and eIF4A1 fractions subjected to a ResourceQ (GE Healthcare) anion exchange column, and eIF4G-MC and eIF4H fractions subjected to Heparin (GE Healthcare) affinity column. Bound protein was eluted with a linear KCl gradient from 100 to 1000 mM KCl. Pooled fractions were further purified by size exclusion chromatography using a Superdex 200 column equilibrated in storage buffer [20 mM Tris/HCl, pH 7.5., 100 mM KCl, 0.1 mM EDTA, 1 % (v/v) glycerol, 1 mM TCEP]. Pooled fractions were concentrated, flash-frozen in liquid nitrogen and stored at −80°C. Protein concentrations were calculated from the absorbance at 280 nm (A280) using extinction coefficients obtained from ExPASy server (**Supplementary Table 5**). All protein preparations showed an A280/A260 ratio ≥ 1.8; for eIF4H the ratio was ≥ 1.5, indicating negligible amounts of contamination by nucleic acids and nucleotides.

### Ribooligonucleotides

RNAs used in this study were purchased from IBA Lifescience and Integrated DNA Technology and are listed in **Supplementary Table 6**.

### Fluorescence-based RNA-binding

For RNA-binding studies 10 – 50 nM FAM-labelled RNAs were incubated with indicated proteins in assay buffer (AB: 20 mM Hepes/KOH, pH 7.5, 100 mM KCl, 1 mM TCEP, 1 % (v/v) DMSO) supplemented with 2 mM AMPPNP/MgCl_2_ in the presence or absence of 100 µM silvestrol (Generon) in 20 µL reactions for 60 min at 25 °C.

For RNA binding in the presence of cofactors (Fig. 1B, Supplementary Fig. 5H-J), 0.5 µM eIF4G or eIF4H were pre-incubated with all components except eIF4A1 for 10 min. Data were normalised using the respective total signal change per condition.

For RNA-release experiments, protein-RNA complexes were formed by incubation of 50 nM FAM-labelled RNA with 5 µM protein in AB ± 100 µM silvestrol + 2 mM ATP in the absence of magnesium. Binding and ATPase-dependent RNA release was initiated by addition of magnesium chloride to a final concentration of 2 mM.

For FRET-based RNA-binding (Fig. 5E and Supplementary Fig. 6L), 50 nM Cy3-Cy5-labelled RNA duplex substrate was incubated alone or with 3 µM eIF4A1 in AB in the presence of 2 mM AMPPNP/MgCl_2_for 60 mins. Competitor AG-RNA was then added to scavenge excess eIF4A1 as indicated in the Figures. Fluorescence-emission spectra in the range 540 – 800 nm were recorded by excitation at 520 nm. Spectra were corrected for Cy5-emission collected from reactions containing only the Cy5-labelled strand. Corrected spectra were then normalised to the maximum Cy3-fluorescence at 565 nm. Relative FRET was calculated according to the equation

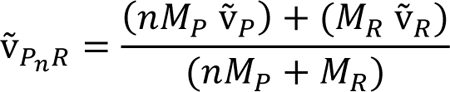

Fluorescence intensities and anisotropy were measured using a Victor X5 (Perkin Elmer) or Spark (Tecan). Dissociation constants and half-lives were obtained from fitting the experimental data to the Hill- and single-exponential decay equation using Prism GraphPad 7 and 8.

### Electrophoretic mobility shift RNA-binding

25 nM Dy680- or Dy780-labelled RNAs were incubated with indicated proteins in AB + 2 mM AMPPNP/MgCl_2_ in the presence and absence of 100 µM silvestrol or 50 µM hippuristanol in 10 µL reactions for 60 min at 25 °C.

In clamping experiments in Fig. 3H, eIF4A1 was preincubated with RNA and silvestrol in AB + 2 mM MgCl_2_ in the absence of nucleotide for 60 min at 25 °C before competitor AG-RNA was added. A final concentration of 2 % (w/v) Ficoll-400 was added to the samples and complexes separated on 6-7% acrylamide-TB gels at 100V for 50 min at room temperature using 0.5xTB as running buffer. When binding of eIF4A1 to the unwinding substrate was analysed, gels were run at 4 °C. Gels were scanned immediately after the run with Odyssey (Licor) or Typhoon-2000 and band intensities quantified using Image Studio Lite. Dissociation constants were obtained from fitting the experimental data to the Hill-equation using Prism GraphPad 7 and 8.

### Analytical gel filtration

eIF4A1 alone or with RNA was incubated for 1 h in AB supplemented with 2 mM AMPPNP/MgCl_2_ -/+ 100 µM silvestrol at room temperature at concentration of 16 µM and 4 µM, respectively, if the protein was in excess; or at 4 µM and 12 µM, respectively, if the RNA was in excess. Samples were loaded onto a S200 increase 3.2/300 (2.4 mL) that was equilibrated in AB + 2 mM MgCl_2_ without AMPPNP. Ovalbumin (45 kDa) and Conalbumin (75 kDa) were used as molecular weight standards.

### Analytical ultracentrifugation

All analytical ultracentrifugation experiments were performed at 50,000 rpm, using a Beckman Optima analytical ultracentrifuge with an An-50Ti rotor at 20°C. Data were recorded using the absorbance optical detection system. For characterisation of the individual protein, sedimentation velocity (SV) scans were recorded at 280 nm in AB ± 2 mM AMPPNP/MgCl_2_ ± 100 µM silvestrol. For characterisation of the individual RNA samples Dy780-(AG)_5_ and 6-FAM-(AG)_10_, SV scans were recorded at 766 nm and 495 nm, respectively, in AB ± 2 mM AMPPNP/MgCl_2_ ± 100 µM silvestrol. For characterisation of the protein in complex with either Dy780-(AG)_5_ or 6-FAM-(AG)_10_, SV scans were recorded at 766 nm or 495 nm, respectively, in either assay buffer ± 2 mM AMPPNP/MgCl_2_ ± 100 µM silvestrol.

The density and viscosity of the buffer was measured experimentally using a DMA 5000M densitometer equipped with a Lovis 200ME viscometer module. The partial specific volume of the protein was calculated using SEDFIT from the amino acid sequence. The partial specific volume of the RNA was calculated using NucProt from the nucleotide sequence. The partial specific volumes of RNA:protein complexes with different stoichiometries were calculated using the equation:

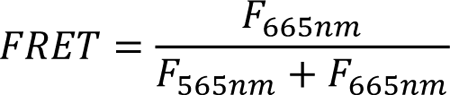

Where *M_P_* and *ṽ_P_* denote the molecular mass and partial specific volume of the protein, respectively, and *M_R_* and *ṽ_R_* denote the molecular mass and partial specific volume of the RNA, respectively. Data were processed using SEDFIT, fitting to the c(s) model.

### Fluorescence lifetime imaging

Fluorescence lifetime measurements in live cells were conducted as described previously (Nobis et al., 2013). Briefly, a Lambert Instruments fluorescence system attached to a Nikon Eclipse TE 2000-U applying LIFA frequency domain method was used. Each sample was excited at 436nm (bandwidth 20 nm). Alongside each experiment, fluorescein in 0.1 M Tris-HCl, pH > 10, was used as a reference of known lifetime of 4 ns. The lifetime of donor for each sample was calculated using the LI-FLIM software (version 1.2.12.30; Lambert Instruments). Presented data (dots) are the average of a technical duplicate per replicate from 9 independent experiments accounting in total for at least 205 cells per condition. For experiments with overexpression of tagged-eIF4A1^DQAD^ as the FRET acceptor (Supplementary Fig. 3J-K), the average of technical duplicate per replicate from 4 independent experiments accounting for 171 cells for donor-eIF4A1wt and 129 cells for donor-eIF4A1wt co-expressed with acceptor-eIF4A1^DQAD^. Representative images are shown with a standard cyan-magenta colour look up table with the limits 3 and 4.2 ns. The scale shown in the images is 50 µm. The line in the dot plot is the mean across all independent replicates.

### Helicase substrates

For fluorescence-based unwinding, overhang, Cy3-reporter and BHQ-quencher strands (Supplementary Table 6) were mixed in a 1.1:1:1 molar ratio in annealing buffer (20 mM Tris-acetate, pH 7.5 and 100 mM KCl). For unwinding gel shifts and FRET-substrates, loading strands and reporters (Supplementary Table 6) were mixed in 1.1:1 ratio in annealing buffer. Reactions were incubated at 85 °C for 15 min and slowly cooled down over 4-5 h in a water bath. Annealed strands were aliquoted and stored at −80 °C.

### Complex formation between eIF4A1 and cofactors

For experiments that included complexes between eIF4A1 and cofactors or combinations thereof, unless otherwise stated proteins have been preincubated in AB in the absence of RNA and nucleotides for at least 60 min before RNA was added.

### Real-time fluorescence-based unwinding

For titrations, 50 nM annealed substrate were incubated with indicated proteins in AB in 18 µL reactions in the presence (clamping conditions) or absence (non-clamping conditions) of 100 µM silvestrol in 384-well plates and incubated for 1 h at 30 °C. Protein dilutions were prepared using storage buffer.

Under scavenging conditions (Fig. 3G-I) 2 µM or indicated concentrations of AG-RNA was added after the pre-incubation step and allowed to scavenge excess eIF4A1 for another 60 min. In pre-clamping experiments i.e., when eIF4A1^wt^ or eIF4A1^DQAD^ were pre-bound to the RNA substrate before addition of the next protein (Fig. 4B), 1 µM indicated eIF4A1 variant was incubated with the RNA substrate in the absence of nucleotide for 1 h before additional eIF4A1 was added to the reaction.

When fractional mixes of eIF4A1^wt^ and eIF4A1^DQAD^ were used, they were first premixed at 50 µM (10x stock) concentration in storage buffer before added to the reaction mixtures. Reactions were started by addition of ATP-MgCl_2_ to a final concentration of 2 mM and fluorescence readings taken in an InfinitePro M200 (Tecan) or Spark (Tecan) with excitation at 535 nm and emission at 575 nm. Data were analysed as described previously (Avanzino et al., 2017; Feoktistova et al., 2013). Data were fitted to a linear equation to yield the initial rate of unwinding as well as the total fraction unwound, respectively. Unless stated otherwise, secondary data were further analysed for the Hill-equation using Prism (GraphPad).

### ATPase assay

ATPase reactions were carried out side-by-side from the same master mix as the fluorescence-based unwinding assays. In separate reactions, NADH (Sigma), phosphoenolpyruvate (Sigma or Alfa Aesar) and lactate dehydrogenase/pyruvate kinase mix (Sigma) were added to unwinding reactions to a final concentration of 2 mM, 2 mM and 1/250 (v/v), respectively. NADH turnover was monitored by measuring absorbance at 340 nm. Obtained absorbance data were converted to the concentration of NADH using condition and machine specific ε (NADH) of 0.62 mM^-1^. ATPase rates were obtained from a linear fit to the experimental data using Prism (GraphPad 7 or 8).

### Unwinding gel shift

All reactions were prepared from the same master mix and split accordingly for the following different conditions. 50 nM annealed substrate was incubated with 3 µM eIF4A1 in AB supplemented with 2 mM MgCl_2_ in 10 µL reactions in the presence or absence of 100 µM silvestrol and incubated for 1 h at room temperature. Under scavenging conditions, a final concentration of 2 µM AG-RNA was added after the preincubation step and allowed to scavenge excess eIF4A1 for another 60 min. Reactions were then started by addition of a final concentration of 2 mM ATP. Reactions were quenched after another 60 min with stop solution (0.5xTBE, 0.2 % (w/v) SDS, 50 mM EDTA, pH 8), or, if RNA-bound complexes were to be resolved, only 2 % (w/v) Ficoll-400 was added. Samples were subjected to gel electrophoresis on discontinuous 10%-acrylamide TB/18%-acrylamide-TBE gels. Gels were run at 200 V at 4 °C and immediately scanned using an Odyssey instrument (LICOR).

### Small-angle X-ray scattering

Samples contained 100 µM eIF4A1 alone, 100 µM eIF4A1 with 30 µM either AG-RNA or AG-overhang substrate to generate multimer eIF4A1-RNA complexes, or 60 µM and 100 µM eIF4A1 with 60 µM AG-RNA or 100 µM CAA-RNA, respectively, to generate monomer complexes in AB supplemented with 2 mM AMPPNP/MgCl_2_ and 100 µM silvestrol. Samples were kept at a concentration of approximately 5mg / ml, frozen in liquid nitrogen, and shipped to Diamond Light Source on dry ice. The protein was applied to a Superdex 200 Increase 3.2 column, at 0.16 ml/min, before being exposed to the X-ray beam, as part of the standard set up at station B21. Data were analysed using ScÅtter version 3.2h. Seventeen *ab initio* models were calculated by DAMMIF (Franke and Svergun, 2009), and average models of these were calculated using DAMAVER and DAMFILT (Volkov and Svergun, 2003). Reported resolution of the space-filled models was calculated using SASRES (Tuukkanen et al., 2016). Superpositions of *ab initio* models were calculated by SUPCOMB (Kozin and Svergun, 2001) or SITUS (Kovacs et al., 2018). Distances shown in Supplementary Table 3 are the mean ± SD based on four individual measurements using PyMOL2. Volumes are the results from data analysis using ScÅtter.

### Reporter mRNA construction

Plasmids containing the desired cDNAs were constructed using annealed oligos (**see Supplementary Table 7**). The sequences were cloned into the pGL3-promoter plasmid (Promega E1761) between the HindIII and NcoI restriction sites, directly upstream of the FLuc open reading frame followed by a 3’UTR and an (A)_49_ sequence. Plasmids were linearised with NsiI located directly downstream of the (A)_49_ sequence and treated with Klenow fragment (NEB M0210S) to generate blunt-ends. RNA was then transcribed from the NsiI-linearised plasmids with the HiScribe™ T7 ARCA mRNA Kit (NEB E2065S) or HiScribe™ T7 mRNA Kit with CleanCap® Reagent AG (NEB E2080S) as per the manufacturer’s instructions. RNAs were purified by acid-phenol chloroform extraction and ethanol precipitation with ammonium acetate and the concentration was quantified spectroscopically and RNA integrity checked by formaldehyde denaturing agarose gel electrophoresis. RNA was stored at −80 °C.

### *In vitro* translation assay

1 ml nuclease-untreated Rabbit Reticulocyte Lysate (Promega L4151) was supplemented with 25 µM haemin, 25 µg/ml creatine kinase, 3 mg/ml creatine phosphate, 50 µg/ml liver tRNAs and 3 mM glucose and aliquoted and stored at −80°C. 50 ng Firefly-luciferase (FLuc) reporter constructs were mixed with storage buffer or storage buffer supplemented with recombinant 4E-BP1 (Sino Biological, 10022-H07E), eIF4A1^wt^ or eIF4A1^E183Q^ (eIF4A1^DQAD^) at room temperature in a volume of 4 µL. Then 11 µl master mix of supplemented untreated Rabbit Reticulocyte Lysate [125 mM KCl, 0.5 mM MgOAc, 20 µM amino acid mix (complete, Promega), 4 U RNaseIn plus Ribonuclease Inhibitor (Promega), 1 mM NaF and 100 µM luciferin (Promega)] to a final volume of 15 µl with water. Final concentrations of recombinant protein were 4 µM 4E-BP1, 16 µM eIF4A1^wt^ or 1 µM eIF4A1^DQAD^. Reactions were prepared in technical duplicates, incubated at 30°C and luciferase activity monitored in real time for 1-2h using a Tecan Spark plate reader. Readings from duplicates were averaged and the maximum translation was extracted as the maximum increase in firefly luciferase (FL) activity over time (slope) (Alekhina et al., 2020). FL activities are shown relative to the FL-activity of the CAA reporter (set to 1) per respective condition.

### RNA Structure-seq2 analysis

To assess changes in RNA structure surrounding polypurine rich sequences, we interrogated our previously published Structure-seq2 data set from MCF7 cells (Waldron et al., 2019), which measured changes in reactivity of RNA structure to dimethyl sulphate (DMS) upon specific inhibition of eIF4A with hippuristanol. DMS modifies non base-paired As and Cs, hence DMS-reactivity is a measure of single-strandedness. The reactivity data are available at the Gene Expression Omnibus (GEO) database accession GSE134865, which can be found at https://www.ncbi.nlm.nih.gov/geo/query/acc.cgi?acc=GSE134865.

To identify all non-overlapping polypurine (R10) sequences in the data set, where R refers to a purine, we made use of the react_composition.py script from the StructureFold2 package of scripts (Tack et al., 2018), which is available from GitHub using the following link https://github.com/StructureFold2/StructureFold2. To exclude A_10_ and G_10_ motifs in our group of polypurine motifs, we only included 10nt 100% R motifs in the analyses with a maximum of 7/10 being purely As or Gs i.e., 100 % R excluding motifs with more than 8 As or Gs. This script outputs the reactivity changes at all motifs and a user defined size either side of the identified motif. Using these data we then filtered the output to the coverage and 5’ end coverage thresholds used previously (Waldron et al., 2019) and picked the most abundant transcript per gene with a 5’UTR length of more than 100 nt, while retaining only those R10 motifs that had at least 3 guanines and 3 adenines. All group sizes are summarised in **Supplementary Table 1**.

For plots Fig. 2F and Supplementary Fig. 2G only those motifs that were positioned at least 50 nt from a UTR/CDS boundary or the 5’ or 3’ end of the transcript were included. This identified 608 R10 motifs in the 5’UTRs of 358 transcripts, 6927 R10 motifs in the CDSs of 1906 transcripts and 2761 R10 motifs in the 3’UTRs of 1303 transcripts. Random motifs were selected using a sliding window analysis (same constraints as R10-analysis, 20nt windows with 10 nt steps) using the react_windows.py script from StructureFold2. The same number of random motifs as R10 motifs were selected from each transcript.

The minimum free energy (MFE, Supplementary Fig. 2I) of predicted folds was calculated by folding the 50 nt windows shown in Fig. 2F centred on the 31:50 downstream window directly downstream of all R10 or random motifs using the batch_fold_RNA.py, which uses RNAstructure (version 6.1) (Reuter and Mathews, 2010) and extracting the metrics with the structure_statistics.py scripts from the StructureFold2 package.

All panels were created using the custom R scripts R10_analysis_1.R and R10_analysis_2.R, which are available at GitHub using the following link https://github.com/Bushell-lab/Structure-seq2-with-hippuristanol-treatment-in-MCF7-cells. The box plot shows the median (centre line), the upper and lower quartile (box limits), the 1.5x interquartile range (whiskers) and in Supplementary Fig. 2I the mean (dot). Outliers (> 1.5x interquartile range) are not shown.

### Calculation of total cellular mRNA concentration

The concentration of mRNAs for a typical HeLa cell was calculated assuming a cell volume of 2425 µm^3^ (BNID: 103725 (Milo et al., 2010) and (Zhao et al., 2008)) and an average mRNA copy number of 300000 per cell (BNID: 104330 (Milo et al., 2010) and (Velculescu et al., 1999)).

### TMT-pulsed SILAC

MCF7 cells were cultivated in SILAC DMEM (Silantes) supplemented with 10 % dialysed FBS (Sigma), 2 mM L-glutamine (Gibco), 0.789 mM Lys-^12^C ^14^N (Lys0) and 0.398 mM Arg-^12^C ^14^N (Arg0), referred to as light-DMEM, for at least 5 doubling times. All isotope-labelled amino acids were purchased from Cambridge Isotope Laboratories with an isotope purity > 99%. For metabolic pulse-labelling, cells were then split using light-DMEM, allowing settling overnight, followed by treatment on the next day with either 150 nM hippuristanol (0.8% DMSO stock) or DMSO control for eight hours in SILAC DMEM (Silantes) supplemented with 10 % dialysed FBS (Sigma), 2 mM L-glutamine (Gibco), 0.789 mM Lys-^13^C ^15^N (Lys8) and 0.398 mM Arg-^13^C ^15^N (Arg10), referred to as heavy-DMEM. Samples were taken immediately at the beginning of treatment (time = 0 h) and after two, four and eight hours after medium swap and treatment. Cells were harvested, washed in PBS and lysed in 6 M urea, 2 M thiourea, 50 mM Tris/HCl pH 8.5, 75 mM NaCl using sonication, and cleared by centrifugation. Supernatants were stored at −80°C. For all time points a biological quadruplet was generated before submission to MS.

25µg protein lysate was reduced with 5 mM DTT, then alkylated in the dark with 50 mM IAA. Samples were then subject to a two-step digestion, firstly with Endoproteinase Lys-C (ratio 1:33 enzyme:lysate) (Promega) for 1 hour at room temperature then with trypsin (ratio 1:33 enzyme:lysate) (Promega) overnight at 37°C. Once digested, peptide samples were labelled with TMT 16plex reagent kit (Thermo Scientific).

400 µg digested sample was fractionated using reverse phase chromatography at pH 10. Solvents A (98% water, 2% ACN) and B (90% ACN, 10% water) were pH adjusted to pH 10 using ammonium hydroxide. Samples were run on an Agilent 1260 Infinity II HPLC. Samples were manually injected using a Rheodyne valve. Once injected the samples were subjected to a two-step gradient, 2-28% Solvent B in 39 mins then 28-46% Solvent B in 13 mins. The column was washed for 8 mins at 100% Solvent B followed by a re-equilibration for 7 mins. Total run time was 76 mins and flow rate was set to 200 µL/min. The samples were collected into 21 fractions.

Peptide samples were run on a Thermo Scientific Orbitrap Lumos mass spectrometer coupled to an EASY-nLC II 1200 chromatography system (Agilent). Samples were loaded onto a 50 cm fused silica emitter (packed in-house with ReproSIL-Pur C18-AQ, 1.9 µm resin) which was heated to 55°C using a column oven (Sonation). Peptides were eluted at a flow rate of 300 nl/min over three optimised two-step gradient methods for fractions 1-7, 8-15 and 16-21. Step one was commenced for 75 mins and step two for 25 mins. For fractionated samples 1-7 the % of solvent B was 3-18% at step one and 30% at step two. For fractions 8-15 the % of B was 5-24% at step one and 38% at step two and for fractions 16-21 the % B was from 7-30% at step one and 47% at step two. Peptides were electrosprayed into the mass spectrometer using a nanoelectropsray ion source (Thermo Scientific). An Active Background Ion Reduction Device (ABIRD, ESI Source Solutions) was used to decrease air contaminants.

Data were acquired using Xcalibur software (Thermo Scientific) in positive mode utilising data-dependent acquisition. Full scan mass (MS1) range was set to 350-1400m/z at 120,000 resolution. Injection time was set to 50 ms with a target value of 5E5 ions. HCD fragmentation was triggered at top speed [3 sec] for MS2 analysis. MS2 injection time was set to 175 ms with a target of 2E5 ions and resolution of 15,000. Ions that have already been selected for MS2 were dynamically excluded for 30 s.

Data were processed following recommendation from Zecha et al. (Zecha et al., 2018) MS raw data were processed using MaxQuant software (Cox and Mann, 2008) version 1.6.14.0 and searched with the Andromeda search engine (Cox et al., 2011) against the Uniprot *Homo Sapiens* database(2018, 95,146 entries). Data were searched with multiplicity set to MS2 level TMT16plex. First and main searches were done with a precursor mass tolerance of 20 ppm for the first search and 4.5 ppm for the main. MS/MS mass tolerance was set to 20 ppm. Minimum peptide length was set to 7 amino acids and trypsin cleavage was selected allowing up to 2 missed cleavage sites. Methionine oxidation and N-terminal acetylation, SILAC Arg10, SILAC Lys8 were selected as variable modifications and Carbimidomethylation as a fixed modification. False discovery rate was set to 1%.

MaxQuant output was processed using Perseus software (Tyanova et al., 2016) version 1.6.15.0. The MaxQuant Evidence.txt file was used to create a new protein groups file. In short, data were culled of contaminant, reverse and unique proteins only peptide identifications before identifying the TMT reporter ion intensities that contain the variable SILAC Arg10 & Lys8 modifications. Identical peptide sequences were combined by median. The data was then exported to R and a script run to combine the ‘TMT reporter intensity corrected’ peptide sequences that belong to the same protein into a ‘protein group’ TMT reporter intensity value (R script available upon request). Protein level data were normalised by LIMMA to account for batch effect differences.

Further data processing and analyses followed recommendation from Zecha et al. (Zecha et al., 2018) To normalise and focus on newly synthesised proteins, which contain the heavy-label (Arg10, Lys8) TMT intensity (I_Lys8,Arg10_), custom R-scripts were used to convert the per-gene intensity data into fraction ‘heavy’ (F_H_) by dividing TMT-intensities from ‘heavy-labelled’ by the sum of light- and heavy-labelled TMT intensities (F_H_ = I_Lys8,Arg10_ / (I_Lys8,Arg10_ + I_Lys0,Arg0_). Considering the limited time points that were collected and to calculate the apparent translation rate of newly synthesised protein, a linearised first order equation for labelling kinetics (single exponential growth) was fitted to the logarithmic data, which yields the apparent translation rate *k* as the slope of the fit (ln F_H_ = *k* * t + offset). To remove poor quality fits, data for the rate *k* for both control and hippuristanol conditions were filtered using a p-value cut-off of 0.1 (two-sided t-test, null-hypothesis of *k* = 0; input: 1337 proteins, p<0.1: 1270, p>0.1: 67). Next, difference and log2-fold change (hippuristanol/DMSO) and associated false discovery rates (FDR) were calculated using standard procedures. Proteins were grouped eIF4A1-dependent or-independent if their FDR of the difference between the apparent translational rate under hippuristanol and DMSO control was smaller than 0.1 or larger than 0.7 respectively. This resulted in 255 eIF4A1-dependent and 244 eIF4A1-independent genes. Motif identification for AG5- and GC5-motif has been done as described under the RNA structure-seq2 section. For Fig. 2G, the same DMS reactivity data as in Fig. 2F was used, and windows were categorised as decrease or increase in RNA structure if their change in DMS reactivity was lower or above 0. In all figures where applicable data is filtered for most abundant transcripts using RNAseq data from our previous study Waldron et al. (Waldron et al., 2019) MFEs in Supplementary Fig. 2E were calculated by averaging the top five folding energies of the most abundant transcript based on mfold prediction (RNA stabilities version 2.3).

### Statistical analysis

If not stated otherwise, n is the number of independent biological replicates of the described experiment and is given in the figure legends. Quantitative experiments including unwinding and ATPase assays and FLIM-FRET were performed in technical duplicates per biological replicate, the average of which was used for downstream analysis. Except for statistical tests based on sequencing data, significance was determined using a two-tailed and unpaired t test. Where applicable p-values were corrected for multiple testing by calculating FDRs. Group sizes are summarised in Supplementary Table 1. Statistical significances are given as the absolute p-values in the figures or figure legends.

## DATA AVAILABILITY

The RNA structure-seq2 datasets analysed were generated previously (Waldron et al., 2019) and are available in the Gene Expression Omnibus (GEO) database accessions GSE134865 and GSE134888 which can be found at https://www.ncbi.nlm.nih.gov/geo/query/acc.cgi?acc=GSE134865 and https://www.ncbi.nlm.nih.gov/geo/query/acc.cgi?acc=GSE134888. All custom scripts and input data are available from GitHub using the following link https://github.com/Bushell-lab/Structure-seq2-with-hippuristanol-treatment-in-MCF7-cells. All StructureFold2 scripts are available from GitHub using the following link https://github.com/StructureFold2/StructureFold2. The TMT-pulsed SILAC mass spectrometry data have been deposited to the ProteomeXchange Consortium via the PRIDE (Perez-Riverol et al., 2022) partner repository with the dataset identifier PXD034343.

## Supporting information

merged supplmentary data, figures and tables

## ACKNOWLEDGEMENTS

We would like to thank the Core Services and Advanced Technologies at the Cancer Research UK Beatson Institute (C596/A17196 and A31287) with particular thanks to the Beatson Advanced Imaging Resource and Molecular Technologies. We thank Diamond Light Source for access to station B21 (BAG allocation MX21657), and the support of N. Cowieson and N. Khunti. We thank S. M. Assman and P. C. Bevilacqua from Penn State University for critical reading of the manuscript and acknowledge their involvement in previous generation of the RNA Structure-seq2 data.

## AUTHOR CONTRIBUTIONS

TS and MB conceived and designed the project. TS, AD, JAW, AW and MB designed experiments. TS and AD performed FA and EMSA RNA-binding assays, ATPase and unwinding assays and analytical gel filtrations. TS cloned protein expression constructs for *in vitro* experiments and designed RNA substrates. AD cloned constructs for FLIM-FRET experiments. TS, AD and JM produced biomass and purified proteins. MG analysed SAXS data with input from TS, AD and DH. TS and JAW performed and analysed *in vitro* translation experiments. JAW performed RNA structure-seq2 related bioinformatics with the support from DCT. AD performed and analysed FLIM-FRET experiments with support from TS, EM and LC. GH and DS performed and analysed analytical ultracentrifugation data with support from TS. TS prepared cell culture samples for pulsed SILAC, KH performed mass spectrometry-related sample processing, mass spectrometry and raw data processing together with GK; TS and KH analysed SILAC data with support from SZ. TS and MB wrote the manuscript with feedback from all authors.

## MATERIAL and CORRESPONDENCE

Further information and requests for resources and reagents should be directed to and will be fulfilled by Tobias Schmidt or Martin Bushell.

