## Supplementary material for "Purine-rich RNA sequences in the 5’UTR site-specifically regulate eIF4A1-unwinding through eIF4A1-multimerisation to facilitate translation": merged supplmentary data, figures and tables

### SUPPLEMENTARY INFORMATION

#### SUPPLEMENTARY RESULTS

##### Cofactors work on distinct eIF4A1-subunits within multimeric eIF4A1- complexes

Cellular eIF4A1 function is believed to be tightly regulated through interactions with its cofactors eIF4H, eIF4B and eIF4G (Andreou and Klostermeier, 2014; Garcia-Garcia et al., 2015; Nielsen et al., 2011; Rogers et al., 2001). We thus asked if the cofactors function upon multimeric eIF4A1. For this, we used full-length eIF4H, which is functionally similar to eIF4B *in vitro* (Rogers et al., 2001), and N-terminally truncated eIF4G-MC (**Supplementary Fig. 5A**), which contains the eIF4A1-binding sites and is eIF4E-independent, thus constitutively active (Feoktistova et al., 2013).

We observed that cofactor activity was exclusively dependent on excess eIF4A1 i.e., the formation of unwinding-competent multimeric complexes (**Supplementary Fig. 5B**) regardless of pre-clamping eIF4A1 to the substrate (**Supplementary Fig. 5C**). We therefore concluded that under scavenging conditions the unwinding substrate and scavenging RNA were not competing for cofactor/eIF4A1-binding. Accordingly, the data showed that cofactors only function upon eIF4A1 under conditions that allow multimerisation. Titrations revealed that cofactor synergy was concentration-dependent (**Supplementary Fig. 5D**), with eIF4H activating eIF4A1 in a 2:1 stoichiometry (Hill-coefficient = 1.7 ± 0.3), while eIF4G-MC only synergised with eIF4A1-eIF4H at submolar concentrations (**Supplementary Fig. 5D**). In contrast to eIF4H, high eIF4G-MC concentrations inhibited eIF4A1 unwinding activity, indicating competition for eIF4A1 subunits (**Supplementary Fig. 5D**). As eIF4G-MC contains two eIF4A1 binding sites (Imataka and Sonenberg, 1997; Lamphear et al., 1995), this also suggested that occupation of both eIF4A1-binding sites is required for an unwinding-competent eIF4G-eIF4A1 interaction.

To understand the competitiveness of eIF4G better, we next examined the effect of the cofactors on unwinding-competent eIF4A1 complexes. eIF4H stimulated eIF4A1-dependent duplex unwinding regardless of the RNA sequence (**Supplementary Fig. 5E and Fig. 1B**), while eIF4G-MC affected maximal unwinding of eIF4A1 only weakly (**Supplementary Fig. 5E and Fig. 1B**) as described before (Andreou and Klostermeier, 2014; Nielsen et al., 2011). Moreover, unwinding of eIF4A1 was generally weaker in the presence of high concentrations of eIF4G-MC as compared to the lower eIF4G concentration (**Supplementary Fig. 5E)**, consistent with a competitive behaviour of eIF4G-MC (**Supplementary Fig. 5D)**. A similar trend was observed when both cofactors were added simultaneously confirming that synergistic activation of eIF4A1 was only optimal under limiting eIF4G and eIF4H concentrations, and that high eIF4G levels inhibited the reaction regardless of eIF4H (**Supplementary Fig. 5F)**. Both cofactors appeared to reduce cooperativity (Hill-coefficients) between eIF4A1 subunits **(Supplementary Fig. 5G)**, while eIF4G-MC at high concentrations also decreased the functional binding affinity of eIF4A1 to the substrate **(Supplementary Fig. 5G)**. Additionally, RNA binding of eIF4A1 in unwinding-competent conditions was limited in the presence of eIF4G-MC, while eIF4H improved eIF4A1's RNA binding (**Supplementary Fig. 5H-I**). Note that, eIF4H and eIF4G-MC bound the substrates at comparable affinities (**Supplementary Fig. 5J**). Collectively, this showed that eIF4H and eIF4G-MC function distinctly upon multimeric eIF4A1 complexes to modulate the unwinding reaction.

To examine the consequence of cofactor binding upon the cooperation between the eIF4A1 subunits mechanistically, we again mixed eIF4A1$^{wt}$ with inactive eIF4A1$^{DQAD}$ at constant total protein concentration but in the presence of the cofactors. eIF4H did not significantly change the cooperation between the eIF4A1 subunits (**Supplementary Fig. 5K**), while eIF4G strongly limited eIF4A1-subunit communication which is reflected by the linear trend of the activities. This suggested that eIF4G, in contrast to eIF4H, now coordinates eIF4A1 subunits and activity potentially by competitive replacement of the loading subunit of eIF4A1, while eIF4H functions mainly upon the unwinding subunits.

As silvestrol clamps the loading subunit to the overhang of the substrate (**Figs. 3F and 3I, Supplementary Fig. 3M**), we hypothesised that eIF4G would not stimulate silvestrol-clamped eIF4A1.

Our results showed that eIF4H and silvestrol synergistically activated eIF4A1, while eIF4G inhibited clamped eIF4A1 (**Supplementary Fig. 5L**) without affecting their interaction (**Supplementary Fig. 5M**).

Collectively, these data clearly demonstrate that eIF4G competes with the eIF4A1 for initial substrate binding to drive a distinct eIF4A1 complex, in which eIF4G replaces the loading subunit of multimeric eIF4A1 complex. In contrast, eIF4H stabilises the eIF4A1-RNA complex and functions upon the unwinding subunits to stimulate unwinding.

#### SUPPLEMENTARY FIGURE LEGENDS

**Supplementary Figure 1. Unwinding by eIF4A1 is stimulated in an RNA sequence-dependent manner *in vitro*. A,** Relative binding affinity of eIF4A1 to 20 nt repeat ssRNA sequences measured as frequency of occurrence in a bind-n-seq study by Iwasaki et al. (Iwasaki et al., 2016) Figure has been created using data from Iwasaki et al. (Iwasaki et al., 2016) Highlighted motifs were used in this study. **B,** Coomassie-stained denaturing SDS-gel after electrophoresis of purified, recombinant eIF4A1. **C**, same as Fig. 1B, but including ATPase under the same conditions. Data are mean ± SEM from repeat experiments, n(ATPase) = 3 for each condition. **D,** Firefly luciferase activity after 60 min translation of reporters shown in Fig. 1C in rabbit reticulocyte lysate in the presence of equimolar or 5-fold excess of 20 nt AG-RNA. AG-RNA in *trans* to the reporter does not stimulate translation of SL-reporter nor affect translation the other reporters. Hence, effects described in Fig. 1D require the AG-repeat motif to be located in *cis* to the 5'UTR of the reporter mRNA. Data are mean from repeat experiments, n = 2. **E**, Indicated regions of the mRNA reporters in red and purple were generated and used as RNA substrates for RNA-binding experiments shown in Supplementary Fig. 1F and unwinding reactions shown in in Supplementary Fig. 1G, respectively. **F**, RNA-binding of eIF4A1 to RNA substrates described in Supplementary Fig. 1E in the presence of AMP-PNP. Data are mean ± sd from three repeat experiments. Binding of eIF4A1 to 5'UTR fragments was about 2-fold stronger if the 166 nt long 5'UTR fragment contained the 20 nt AG repeat sequence as shown in Supplementary Fig. 1E. **G**, unwinding of eIF4A1 using a 5' overhang 24bp substrate with 60 nt CAA-overhang with and without a 20nt AG-repeat 30 nt upstream the duplex region. The design of the substrate reflects the organisation of the 5'UTR fragment indicated in purple in Supplementary Fig. 1E. Data are mean ± sd from two repeat experiments Unwinding by eIF4A1 was enhanced if the RNA substrate contained the AG-repeat sequence, showing that the 20 nt AG-repeat in the 5'UTR recruits eIF4A1 and stimulates its unwinding activity. **H**, Progress curves of $eIF4A1^{wt}$ and $eIF4A1^{DQAD}$ on AG-overhang substrate confirming the unwinding-inactive state of $eIF4A1^{DQAD}$. Data are mean from a technical duplicate.

**Supplementary Figure 2. eIF4A1 shows RNA sequence-dependent unwinding and stimulation of translation in cells. A,** Pearson correlation coefficients between replicates of the TMT-pulsed SILAC from main figure 2, **B,** Histogram showing binned p-values (bins = 0.05) of the derived translation rates from the linear fit of the experimental TMT pulsed SILAC data from control DMSO and hippuristanol conditions. p-values were calculated from a F-test against k(slope) = 0. **C**, Scatter plot of translation rates under control (DMSO) or hippuristanol conditions grouped by p-value cut-off. Genes coloured in green (p-values <0.1, from fitting process supplementary Fig. 2B) were used for downstream analyses. n(green) = 1270, n(purple) = 67. **D**, Density plot of translation rates from DMSO control or hippuristanol conditions. In agreement with hippuristanol being a translational inhibitor, translation rates are globally downregulated upon eIF4A1 inhibition with hippuristanol. The p-value was calculated by a paired, two-sided Wilcoxon test. **E**, 5'UTR features of eIF4A1-dependent and –independent 5'UTRs mRNAs filtered for most abundant transcripts. 5'UTRs were folded and their mean free energy ($\Delta G$) calculated using mfold server (Markham and Zuker, 2008; Zuker, 2003). Thermodynamic folding stabilities of the 5'UTRs of eIF4A1-dependent and –independent mRNAs are comparable. The p-values were calculated by unpaired, two-sided Wilcoxon tests. **F**, Box plots of the change in DMS-reactivity following eIF4A1 inhibition of the 5'UTR, CDS and 3'UTR of eIF4A1-dependent and –independent genes identified in the TMT-pSILAC. This shows that eIF4A1-dependent and -independent mRNAs experience a comparable change in RNA structure after eIF4A1-inhibition. This is in agreement with our previous report, that eIF4A1-inhibtion does not lead to global changes in RNA structure, but localized RNA structure (Waldron et al., 2019). The p-values were calculated by unpaired, two-sided Wilcoxon tests. **G**, upper panel, Schematic presentation of the conducted sliding window analysis. The DMS-reactivity of 20 nt sliding windows with 10 nt step size up- and downstream of 10 nt polypurine (AG5) or random motifs (control) have been compared pairwise. Lower panel, box plot of the average change in DMS-reactivity (ΔDMS reactivity) following eIF4A inhibition of all of 20 nt sliding windows down- and upstream of all 10 nt AG5 motifs in 5'UTRs. The Δ reactivity is calculated by subtracting the reactivity under control conditions from the reactivity under hippuristanol conditions, based on previous RNA Structure-seq2 data in MCF7 cells (Waldron et al., 2019). Therefore, a negative Δ reactivity indicates decreased reactivity following eIF4A inhibition, which is consistent with increased structure and *vice versa*. Changes in RNA structure around AG-motifs are most pronounced in a 20 nt window, 31 nt downstream of the AG motif which is also shown in Fig. 2F. The p-value was calculated by a paired, two-sided Wilcoxon test. **H**, Histogram depicting the binned position of all AG5 motifs in all 5'UTRs more than 100 nt in length from the most abundant transcript per gene in MCF7 cells based on Structure-seq2 data (Waldron et al., 2019). The dotted line depicts the median, which along with the bars indicates that R10 motifs are spread evenly throughout 5'UTRs. n = 1916 motifs from 1186 transcripts. **I**, Violin plots comparing the minimum free energy of the same 5'UTR windows shown in Fig. 2F. Dot is the mean; box plot shows the median. The p-value was calculated by a paired, two-sided Wilcoxon test. **J,** Box plots of the log2-fold change in translation rate following eIF4A1 inhibition of mRNAs containing 5'UTR AG5 motifs (or random control motifs) that show changes in local RNA structure 50 nt upstream the motif (same windows as Fig. 2F). Note, that Fig. 2G shows corresponding downstream RNA regions as defined in Fig 2F. The p-value was calculated by a paired, two-sided Wilcoxon test.

**Supplementary Figure 3. RNA sequence-specific unwinding by eIF4A1 is performed by a multimeric RNA-eIF4A1 complex. A-B,** Unwinding activity of eIF4A1 on a 20 nt AG-overhang substrate with 11 bp or 24 bp duplex region at indicated substrate concentrations. In all cases, a non-hyperbolic but sigmoidal shape of the curve is observed suggesting that independent of the duplex length unwinding by eIF4A1 involves more than one copy of the protein; data are mean (of technical duplicates) from three repeat experiments, n = 3. The dashed lines represent a hypothetical model with h = 1 in which eIF4A1 follows non-sigmoidal kinetics for a monomeric enzyme using the kinetic parameters from fitting the Hill-equation to the experimental data. **C,** ATPase activity, n(repeat experiments) = 3, of eIF4A1 on AG- and CAA-overhang substrate. The Hill-equation was fitted to the data. In contrast to the unwinding reactions shown in main Fig. 3A, the ATPase activity of eIF4A1 shows a non-cooperative behaviour. **D**, Representative electrophoretic mobility shift assays (EMSA) of 20 nt AG- or CAA-RNA at increasing eIF4A1 concentrations. n = 3 repeat experiments. **E,** Representative electrophoretic mobility shift assays of eIF4A1-binding to 25 nM AG- and CAA-RNA in the presence of silvestrol at increasing concentrations of eIF4A1. Multimeric eIF4A1-RNA complexes are detectable only with the AG-RNA; n = 3 repeat experiments. **F**, Analytical gel filtrations of eIF4A1-AG-RNA complexes at saturating protein (16 µM eIF4A1, 4 µM RNA) or RNA concentrations (4 µM eIF4A1, 12 µM RNA) in the presence or absence of silvestrol. Reference protein: Ovalbumin – 45 kDa at 1.6 mL, Conalbumin – 76 kDa at 1.5 mL, free RNA at 2.05 mL (not shown). **G,** Analytical ultracentrifugation of eIF4A1-AG-RNA complexes. eIF4A1 at 8 µM and RNA at 2 (eIF4A1>AG-RNA) or 8 µM (eIF4A1=AG-RNA). Data are summarised in Supplementary Table 2. This confirms results from the gel filtrations that eIF4A1-multimerisatoin is only induced upon RNA-binding. In addition, multimerisation is observed even on the short 10 nt AG-RNA suggesting that in multimeric eIF4A1 only one eIF4A1 molecule is in tight contact with the RNA as shown in a recent crystal structure of eIF4A1 in complex with the same 10 nt AG-RNA in the presence of rocaglamide A, a silvestrol derivate (Iwasaki et al., 2019). Modelling of the data (Supplementary Table 2) suggests a 3:1 stoichiometry of eIF4A1:AG-RNA in the multimeric complexes. **H,** EMSA comparing multimerisation of eIF4A1$^{wt}$ and mTurquoise-eIF4A1 upon binding to AG-RNA. The N-terminal fusion of mTurquoise to eIF4A1 does not affect its ability to form multimeric complexes. **I,** Western blots corresponding to Fig. 3D-E showing expression levels of overexpressed mTurquoise- (mT) and mCitrine-tagged (mC) eIF4A1 (anti-GFP blot) versus endogenous eIF4A1 (anti-eIF4A1 blot) expression. Vinculin is used as the loading control. uncropped blots are shown. **J**, FLIM-FRET images and quantification of co-transfection of mTurquoise-eIF4A1-wildtype and mCitrine-eIF4A1$^{DQAD}$; n = 4 independent repeat experiments. Total cells included all replicates (DQAD) = 129, (wt) = 171; p-value was calculated by an unpaired, two-tailed t-test. The scale is 50 µm. Cotransfection of tagged eIF4A1$^{wt}$ and eIF4A1$^{DQAD}$ yielded a similar change in fluorescence lifetime as with eIF4A1$^{wt}$ only (see Fig. 3E) indicating that RNA-binding capacity of at least one of the subunits in multimeric eIF4A1 is not essential for multimerisation. Hence a direct eIF4A1-eIF4A1 interaction induced by binding of one eIF4A1 to RNA is suggested. **K**, AG-RNA binding of eIF4A1$^{wt}$ and eIF4A1$^{DQAD}$ confirming the RNA-binding deficiency of eIF4A1$^{DQAD}$. **L,** ATPase-dependent RNA release of eIF4A1-AG-RNA complexes ± silvestrol. 4 µM protein were pre-incubated for 60 min with 1 µM AG-RNA in the presence or absence of 100 µM silvestrol together with ATP in the absence of magnesium. Then magnesium chloride was added to a final concentration of 2 mM and after 10 min incubation real-time fluorescence was recorded. Data are the mean of a technical duplicate. AG-RNA is not released in the presence of silvestrol while in the absence of silvestrol the RNA is released as a result of ATP hydrolysis. This shows that silvestrol clamps eIF4A1 to the AG-RNA in an ATP-independent manner as observed previously (Iwasaki et al., 2016). **M**, Unwinding activity of clamped and non-clamped eIF4A1 on 20 nt AG-overhang substrate. The Hill-equation was fitted to the data. Data are mean (of technical duplicates) from three repeat experiments. The back line models the experimental data assuming a non-cooperative ($h = 1$) mechanism. This shows that clamping eIF4A1 to the overhang stimulates unwinding when eIF4A1 is in excess over the RNA and that even under clamping conditions eIF4A1 unwinds RNA as a multimeric enzyme because $h > 1$.

**Supplementary Figure 4. Division of functions in multimeric eIF4A1. A,** EMSA of AG-RNA binding and **B**, progress curve of unwinding of eIF4A1wt and eIF4A1$^{DQAD}$ both in the presence of silvestrol. A technical duplicate is shown. eIF4A1$^{DQAD}$ is catalytically inactive while it binds AG-RNA at similar affinity as eIF4A1wt in the presence of silvestrol. **C**, Representative progress curves (of three replicates) of AG-RNA release of eIF4A1$^{wt}$ and eIF4A1$^{DQAD}$ in the presence of ATP and silvestrol -/+ unlabelled competitor AG-RNA under conditions equivalent to the experiment shown in Figure 4B. The mean ± sd of a technical duplicate is shown. This shows that RNA-complexes of both wildtype and variant eIF4A1 display very similar stabilities under conditions equivalent to those in Figure 4B, arguing against eIF4A1$^{wt}$ replacing overhang-bound eIF4A1$^{DQAD}$. Thus, variant and wildtype protein cooperate to achieve unwinding as shown in Fig. 4B **D**, Progress curves of unwinding by indicated mixes of eIF4A1$^{wt}$ and eIF4A1$^{DQAD}$ at a total concentration of 5 µM following the reaction scheme shown in Fig 4B. A technical duplicate is shown. **E,** ATPase activities of unwinding competent multimeric eIF4A1 complexes comparing eIF4A1$^{wt}$ complexes with ones in which inactive eIF4A1$^{DQAD}$ was pre-clamped to the overhang region. Data are mean ± sd from three repeat experiments; n = 3. The result shows that the ATPase is stronger when eIF4A1$^{wt}$ is bound to the overhang. As the chimeric eIF4A1wt-eIF4A1$^{DQAD}$ complex also unwinds RNA efficiently (Fig. 4B), this suggests that the ATPase activity of actively unwinding eIF4A1 is lower than the eIF4A1-loading subunits that are bound to the overhang region.

**Supplementary Figure 5. eIF4A1-cofactors function upon distinct eIF4A1 subunits within multimeric eIF4A1**. **A,** Coomassie-stained denaturing SDS-gel after electrophoresis of recombinant eIF4H and eIF4G-MC. **B,** Progress curves of 5 µM eIF4A1 on AG-overhang substrate in the presence of 500 nM eIF4H or 500 nM eIF4H and 500 nM eIF4G under excess or scavenged eIF4A1 conditions (See Fig. 3F). Data points are the mean of a technical duplicate ± sd. Unwinding by eIF4A1 is enhanced under excess eIF4A1 conditions even in the presence of the cofactors. **C,** Progress curves of 5 µM eIF4A1 on AG-overhang substrate in the presence of 5 µM eIF4H in the presence or absence of silvestrol. Data points are the mean of a technical duplicate ± sd. Regardless of clamping eIF4A1 to the substrate, unwinding is enhanced when excess eIF4A1 is present, suggesting that under 'scavenging' conditions RNA substrate and cofactor binding do not compete for eIF4A1. **D,** Unwinding activity of 2 µM eIF4A1 (blue) or 2 µM eIF4A1-eIF4H (1:1, green) at increasing concentrations of eIF4H (blue) or eIF4G (green), respectively; data are mean (of technical duplicates) from three repeat experiments; n = 3. The Hill-equation was fitted to the eIF4H-data and an equation describing substrate inhibition was fitted to the eIF4G-data (lines). **E,** Unwinding activity of eIF4A1 on AG- (blue) or CAA-overhang (purple) substrate in the presence of indicated concentrations of eIF4H (left panel) or eIF4G (right panel). Data are mean (of technical duplicates) from three repeat experiments; n=3. The Hill-equation was fitted to the data (lines). **F,** Unwinding activity of eIF4A1 on AG-overhang substrate in the presence of 500 nM (left panel) or 5 µM eIF4H (right panel) together with indicated concentrations of eIF4G. Data are mean (of technical duplicates) from three repeat experiments, n =3. The Hill-equation was fitted to the data (lines). The bottom panel show the change in unwinding activity as compared to in the absence of eIF4G. This shows that only when both cofactors are limiting synergistic stimulation of eIF4A1 by the cofactors is optimal. **G**, Hill-coefficient and functional binding affinities as derived from the fits to the experimental data shown in Supplementary Figure 5E. Bars show result and errors of the fit. Both cofactors lower the Hill-coefficient of unwinding by eIF4A1. Only eIF4G also lowers the apparent functional binding affinity indicating competition with eIF4A1 for substrate binding. **H**, RNA binding of eIF4A1 alone versus RNA-binding of eIF4A1 in the presence of eIF4G-MC or **I**, the presence of eIF4H; data are mean ± sd from three repeat experiments; n = 3. eIF4G reduces eIF4A1-RNA binding while eIF4H stimulates eIF4A1 RNA-binding activity. This confirms that eIF4G competes with eIF4A1 for RNA-binding. **J,** RNA-binding affinities of eIF4G and eIF4H to indicated RNAs. Data are mean ± SD from three repeat experiments, n =3. **K,** Inhibition of unwinding of eIF4A1$^{wt}$ by fractional mixes with eIF4A1$^{DQAD}$ alone (black) in the presence of 5 µM eIF4H (blue) or 2 µM eIF4H with 1 µM eIF4G (green) corresponding to the points of maximum stimulation as shown in Supplementary Fig. 5D. Activity data of each replicate is plotted relative to non-inhibited eIF4A1 (no added eIF4A1$^{DQAD}$); data are mean (of technical duplicates) from three repeat experiments, n = 3. The line indicates the behaviour of a monomeric or multimeric enzyme without subunit-cooperativity. In the presence of eIF4H the cooperation between eIF4A1 subunits is maintained, while in the presence of eIF4G eIF4A1-subunit cooperativity is lost. As unwinding is under all conditions still performed in a cooperative manner (Supplementary Fig 5E-F), this suggests that eIF4G is coordinating the activity of the eIF4A1 unwinding subunits. This might be achieved by eIF4G replacing the loading subunit of multimeric eIF4A1. **L,** Unwinding activity of clamped eIF4A1 on AG-overhang substrate alone (black) and in the presence of 5 µM eIF4H (blue) or eIF4G (green). Data are mean (of technical duplicates) from three repeat experiments, n = 3. The Hill-equation was fitted to the data (lines). This shows that eIF4H stimulates clamped eIF4A1 while eIF4G is inhibitory to the activity of clamped eIF4A1 suggesting that eIF4H functions upon the unwinding-subunits of eIF4A1 while eIF4G operates upon or replaces the loading-subunits of multimeric eIF4A1. **M,** Coomassie-stained SDS-gel of coimmunoprecipitated recombinant mTurquoise-eIF4A1 and eIF4G-MC using an anti-eIF4A1 antibody in the presence of indicated eIF4A1 inhibitors. None of the eIF4A1-inhibitors disrupt the eIF4A1-eIF4G interaction.

**Supplementary Figure 6. The architecture of eIF4A1-loading complexes. A,** Superposition of the envelope model of apo-eIF4A1 (grey) derived from our SAXS data with the crystal-structure of *S. cerevisiae* eIF4A (yellow) representative of an half-open conformation of eIF4A (PDBid 2vso (Schutz et al., 2008), $\chi^2$ = 10), and to the crystal structure of eIF4A1 in complex with a 10nt AG-RNA which is representative of the closed conformation (PDBid 5zc9 (Iwasaki et al., 2019), $\chi^2$ = 163). This suggests that the derived SAXS-model for solution apo-eIF4A1 is more representative of an open and more dynamic conformation than a closed and compact one. **B,** Dimensionless Kratky and **C** Guinier plots of SAXS data corresponding to apo-eIF4A1 and monomeric eIF4A1 bound to AG or CAA-RNA as shown in Fig. 5A. The Kratky plot demonstrates that, while all proteins are in a folded and globular state (bell shape), the eIF4A1-AG-RNA complex is more compact than the CAA-RNA complex and the apo-eIF4A1 (broader curve of CAA-eIF4A1 and apo-eIF4A1 *vs* AG-eIF4A1). The slope of the linear fit in the Guinier plot is used to calculate the radius of gyration. The good quality of the linear fit to the data in the Guinier plot shows negligible aggregation in the samples. **D,** Linear free energy relationships (LFER) between the association constant ($K_A = 1/K_D$) of eIF4A1 to AG- and CAA-RNA and increasing potassium chloride concentrations in the presence or absence of silvestrol. Data are mean ± sd from at least two repeat experiments ($n \geq 2$). The slopes of the linear fits are measures for the relative contribution of ionic and non-ionic interactions in the binding process (Schmidt et al., 2016). Our results show that the addition of silvestrol reduced the dependency of eIF4A1 binding to AG-RNA on the salt concentration. Hence, in the presence of silvestrol, there are additional non-ionic interactions contributing to the RNA-protein interface than in the absence of silvestrol. This is in agreement with the recent crystal structure of eIF4A1 and a 10 nt AG-repeat RNA in the presence of the silvestrol-derivate rocaglamide (Iwasaki et al., 2019) that reveals hydrophobic base stacking interactions between the RNA and the eIF4A1-rocA complex bridged by rocA. The LFER slopes of eIF4A1 binding to AG and CAA RNA are similar in the presence of silvestrol suggesting a comparable contribution of ionic interactions between these two eIF4A1-RNA complexes. As the $K_A$ are different, this accordingly suggests that there is a higher fraction of non-ionic interactions present in the eIF4A1-AG-RNA interface than in the eIF4A1-CA-RNA interface. Note, that we could not measure a LFER for CA-RNA-eIF4A1 in the absence of silvestrol accurately as affinities were very weak at increased salt concentrations. **E,** Dimensionless Kratky and **F** Guinier of SAXS data corresponding to monomeric and multimeric eIF4A1 as shown in Fig 5B. The good quality of the linear fit to the data in the Guinier plot shows negligible aggregation in the samples. **G**, Superposition of the SAXS models corresponding to the monomeric (blue) and multimeric (purple) eIF4A1-AG-RNA complexes. **H,** Dimensionless Kratky and **I** Guinier of SAXS data corresponding to multimeric eIF4A1 bound to AG-ssRNA or AG-overhang substrate as shown in Fig 5D. The Kratky plot of the eIF4A1-overhang substrate complex shows that is has a more extended shape than the eIF4A1-AG-RNA complex and indicates that it has two diffracting domains as suggested by the shoulder in the curve. The good quality of the linear fit to the data in the Guinier plot shows negligible aggregation in the samples. **J**, A 24 bp dsRNA (yellow, extracted from PDBid 2L3J) was manually fitted into the experimental SAXS envelope of multimeric eIF4A1 bound to the overhang of a 24 bps duplex substrate using PyMOL2. This suggests that multimeric eIF4A1 not only binds the single stranded overhang region of the RNA substrate but also, in addition, is directly located at the ssRNA-duplex fork and may cover ~ 5 bps of the duplex region which depicts the hypothetical priming mechanism of unwinding. **K,** Representative (of four repeat experiments) fluorescence-emission spectra (excitation at 520 nm) of FRET-labelled AG-overhang substrate in its unbound/free state (black) or bound by multimeric eIF4A1 (purple). **L,** Relative FRET efficiency of AG-overhang substrate when bound to eIF4A1 derived from fluorescence emission spectra (example shown in Supplementary Figure 6K); data from four repeat experiments; n = 4. Binding of multimeric eIF4A1 to AG-overhang region induces a specific conformational change in the ssRNA-region of the substrate that is different from eIF4A1-monomer binding. Scavenger RNA was added to the multimeric eIF4A1 state (see Fig. 3F-G) to monitor conformational changes in the RNA when multimeric eIF4A1 transitions into its monomeric state (blue). The different relative FRET efficiencies indicate that free overhang as well as monomeric and multimeric eIF4A1-bound overhang adopt different conformations. The FRET at 6 µM scavenger is shown in main Figure 5E.

#### SUPPLEMENTARY TABLES

**Supplementary Table 1. Group sizes of TMT-pSILAC and RNA Structure-seq2 analyses**

| | | Group size (number of transcripts) | motifs |
|---|---|---|---|
| TMT-pSILAC | eIF4A1-dependent | 255 | |
| | eIF4A1-independent | 244 | |
| | total DMSO unfiltered / filtered | 1337 / 1270 | |
| | total hippuristanol unfiltered / filtered | 1337 / 1270 | |
| RNA structure-seq2 | 5'UTR | 358 | 608 |
| | CDS | 1906 | 6927 |
| | 3'UTR | 1303 | 2761 |
| | AG location | | 1617 |

**Supplementary Table 2. Model and experimental parameters of eIF4A1-RNA complexes calculated from analytical ultracentrifugation**

| experimental parameters | | | | | | | Expected parameters | |
|---|---|---|---|---|---|---|---|---|
| Sample | Modelled as AG-RNA:eIF4A1 | Peak ③ MW (kDa) | Peak ③ Sed. Co. (S) | Peak ④ MW (kDa) | Peak ④ Sed. Co. (S) | Frictional ratio | AG-RNA:eIF4A1 | Expected MW (kDa) |
| **eIF4A1 > AG-RNA** | 1:1 | - | - | 107.0 | 6.49 | 1.30 | 1:1 | 53.4 |
| | 1:2 | | | 115.0 | | | 1:2 | 99.4 |
| | 1:3 | | | 118.0 | | | 1:3 | 145.4 |
| | 1:4 | | | 120.0 | | | 1:4 | 191.4 |
| **eIF4A1 = AG-RNA** | 1:1 | 53.8 | 4.12 | - | - | 1.269 | | |
| | 1:2 | 57.5 | | | | | | |
| | 1:3 | 58.9 | | | | | | |
| | 1:4 | 59.7 | | | | | | |

Notes: This shows that when eIF4A1 > AG-RNA the experimental data is best described by a stoichiometry of eIF4A1:RNA of 1:2 or 1:3 (blue), as 1:4 would is far off the expected molecular weight of 191.4 kDa. When eIF4A1 = AG-RNA the experimental data is best described by a stoichiometry of 1:1 (red).

**Supplementary Table 3. Model parameters of eIF4A1-AG-RNA envelopes**

| | | $R_g$ (Å) | $D_{max}$ (Å)* | $V_C$ (a.u.) |
|---|---|---|---|---|
| apo-eIF4A1 | experimental | 30.8 ± 1.1 | 99 ± 5.9 | 223 ± 65 |
| monomer | experimental | 27.3 ± 0.5 | 97.0 ± 7.0 | 175 ± 55 |
| multimer | experimental | 34.6 ± 1.46 | 118.7 ± 6.6 | 693 ± 231 |
| | linear model** | 52.4 | 186.4 | |
| | non-linear model** | 27.4 | 89.4 | |

* $D_{max}$ is derived from the P(r) distribution of the experimental data

** without RNA; a.u. – arbitrary units

**Supplementary Table 4. primers for generation of constructs**

| Name | Sequence 5' to 3' | identifier |
|---|---|---|
| eIF4A1_BsaI_Sumo_fw | TTG GTC TCA TGG TTC TGC GAG CCA GGA TTC C | TS3 |
| eIF4A1_NotI_stop_rv | AAT AGC GGC CGC TCA GAT GAG GTC AGC AAC ATT G | TS4 |
| eIF4G_MC_blunt | GGC CGG ACA ACC CTT AGC ACC CGT G | TS9 |
| eIF4G_NotI_stop_rv | AAT AGC GGC CGC TCA GTT GTG GTC AGA CTC CTC | TS10 |
| eIF4H_BsaI_Sumo_fw | TTG GTC TCA TGG TGC GGA CTT CGA CAC CTA CG | TS15 |
| eIF4H_NotI_stop_rv | AAT AGC GGC CGC TCA TTC TTG CTC CTT TTG AAC GAC | TS16 |
| eIF4A1_DEAD_DQAD_fw | GTT TGT ACT GGA TCA GGC TGA CGA AAT GTT AAG | TS23 |
| eIF4A1_DEAD_DQAD_rv | CTT AAC ATT TCG TCA GCC TGA TCC AGT ACA AAC | TS24 |
| A1_HindIII_fw | GTT GAA GCT TCA TCT GCG AGC CAG GAT TCC | TS64 |
| A1_BamHI_rev | GAA TAG GAT CCT CAG ATG AGG TCA GCA ACA TTG | TS65 |
| Citrine/turquiose-toSumo_fw | TTG GTC TCA TGG TGT GAG CAA GGG CGA G | TS74 |
| pETSUMO_rev_bl | ACC ACC AAT CTG TTC TCT GTG AGC CTC AAT AAT ATC | TS1 |
| pETSUMO_MCS_fw | AGA GAC CTC AGG ATC CAA GCT TGC GG | TS2 |

**Supplementary Table 5. Theoretical protein parameters**

| protein | kDa | ε280 | ε260 | ε280/ ε260 |
|---|---|---|---|---|
| eIF4A1 | 46 | 34630 | 20641 | 1.68 |
| eIF4G-MC | 95 | 69495 | 45702 | 1.52 |
| eIF4H | 25 | 8940 | 8122 | 1.10 |
| $eIF4A1^{DQAD}$ | 46 | 34630 | 20641 | 1.68 |

**Supplementary Table 6. RNAs used in this study**

| Name | Sequence 5' to 3' | Length (nt) | Label and position |
|---|---|---|---|
| AG-RNA | AGAGAGAGAGAGAGAGAGAG | 20 | 5-FAM-EX or 5' Dy780 |
| 10 nt AG-RNA | AGAGAGAGAG | 10 | 5-FAM-EX |
| scavenger RNA | AGAGAGAGAGAGAGAGAGAG | 20 | none |
| CAA-RNA | CAACAACAACAACAACAACA | 20 | 5-FAM-EX or 5' Dy780 |
| AGUG | AGUGAGUGAGUGAGUGAGUG | 20 | 5-FAM-EX |
| UGUU | UGUUUGUUUGUUUGUUUGUU | 20 | 5-FAM-EX |
| UCUC | UCUCUCUCUCUCUCUCUCUC | 20 | 5-FAM-EX |
| Unwinding substrate-20 nt overhang 24 bp duplex | $N_{20}$GAAAAAAUUAAAAAAUUAAAAAAC | 44 | 3' BHQ2 |
| Reporter 24 bp duplex | GUUUUUUAAUUUUUUAAUUUUUUC | 24 | 5' Cy3 |
| AG-overhang (11 bp duplex) | $N_{20}$GACACAACUAC | 31 | 3' BHQ2 |
| Reporter 11 bp | GUA GUU GUG UC | 11 | 5' Cy3 |
| FRET-loading | AGAGAGAGAGAGAGAGAGAGGAAAAAAUUAAAAAAUUAAAAAAC | 44 | 5' Cy3 |
| FRET-reporter | GUUUUUUAAUUUUUUAAUUUUUUC | 24 | 5' Cy5 |
| EMSA-loading | AGAGAGAGAGAGAGAGAGAGGAAAAAAUUAAAAAAUUAAAAAAC | 44 | 5' Dy780 |
| EMSA-reporter | GUUUUUUAAUUUUUUAAUUUUUUC | 24 | 5' Dy680 |

**Supplementary Table 7. DNA primers used to generate reporter mRNAs**

| Name | Sequence 5' to 3' | reporters: CAA | CAA-AG | SL | SL-AG |
|---|---|---|---|---|---|
| CAA_f1 | AGCTTTAATACGACTCACTATAGGGCAAGAACAACAACAACAACAACAACAACAACAACAACAACAACAACAACAACAACAACAACAACAACAACAACAACAACAACAACAACAACAACAACA | X | | X | |
| CAA_f2 | ACAACAACAACAACAACAACAACAACAACAACAACAACAACAACAACAACAACAACAACAACAACAACAC | X | X | | |
| CAA_r1 | GTTGTTGTTGTTGTTGTTGTTGTTGTTGTTGTTGTTGTTGTTCTTGCCCTATAGTGAGTCGTATTAA | X | X | X | X |
| CAA_r2 | CATGGTGTTGTTGTTGTTGTTGTTGTTGTTGTTGTTGTTGTTGTTGTTGTTGTTGTTGTTGTTGTTGTTGTTGTTGTTGTTGTTGTTGTTGTTGTTGTTGTTGTTGTTGTTGTTGTTGTT | X | | | |
| SL_f2 | ACGCGGCGGCGGCGAGAGCAGCAGCAGCAACAACAACAAGCGGCGGCGGCGAGAGCAGCAGCAGCAC | | | X | |
| SL_r2 | CATGGTGCTGCTGCTGCTCTCGCCGCCGCCGCTTGTTGTTGTTGCTGCTGCTGCTCTCGCCGCCGCCGCGTTGTTGTTGTTGTTGTTGTTGTTGTTGTTGTTGTTGTTGTTGTTGTTGTT | | | X | |
| AG_f1 | AGCTTTAATACGACTCACTATAGGGCAAGAACAACAACAACAACAACAACAACAACAACAACAACAACAACAAGAGAGAGAGAGAGAGAGAGAACAACAACAACAACAACAACAACAACA | | X | | X |
| AG_r2 | CATGGTGTTGTTGTTGTTGTTGTTGTTGTTGTTGTTGTTGTTGTTGTTGTTGTTGTTGTTGTTGTTGTTGTTGTTGTTGTTGTTGTTGTTGTTGTTGTTGTTCTCTCTCTCTCTCTCTCTT | | X | | |
| AG_SL_f2 | ACGCGGCGGCGGCGAGAGCAGCAGCAGCAACAACAACAAGCGGCGGCGGCGAGAGCAGCAGCAGCAC | | | | X |
| AG_SL_r2 | CATGGTGCTGCTGCTGCTCTCGCCGCCGCCGCTTGTTGTTGTTGCTGCTGCTGCTCTCGCCGCCGCCGCGTTGTTGTTGTTGTTGTTGTTGTTGTTGTTCTCTCTCTCTCTCTCTCTT | | | | X |

### Supplementary Figure 1

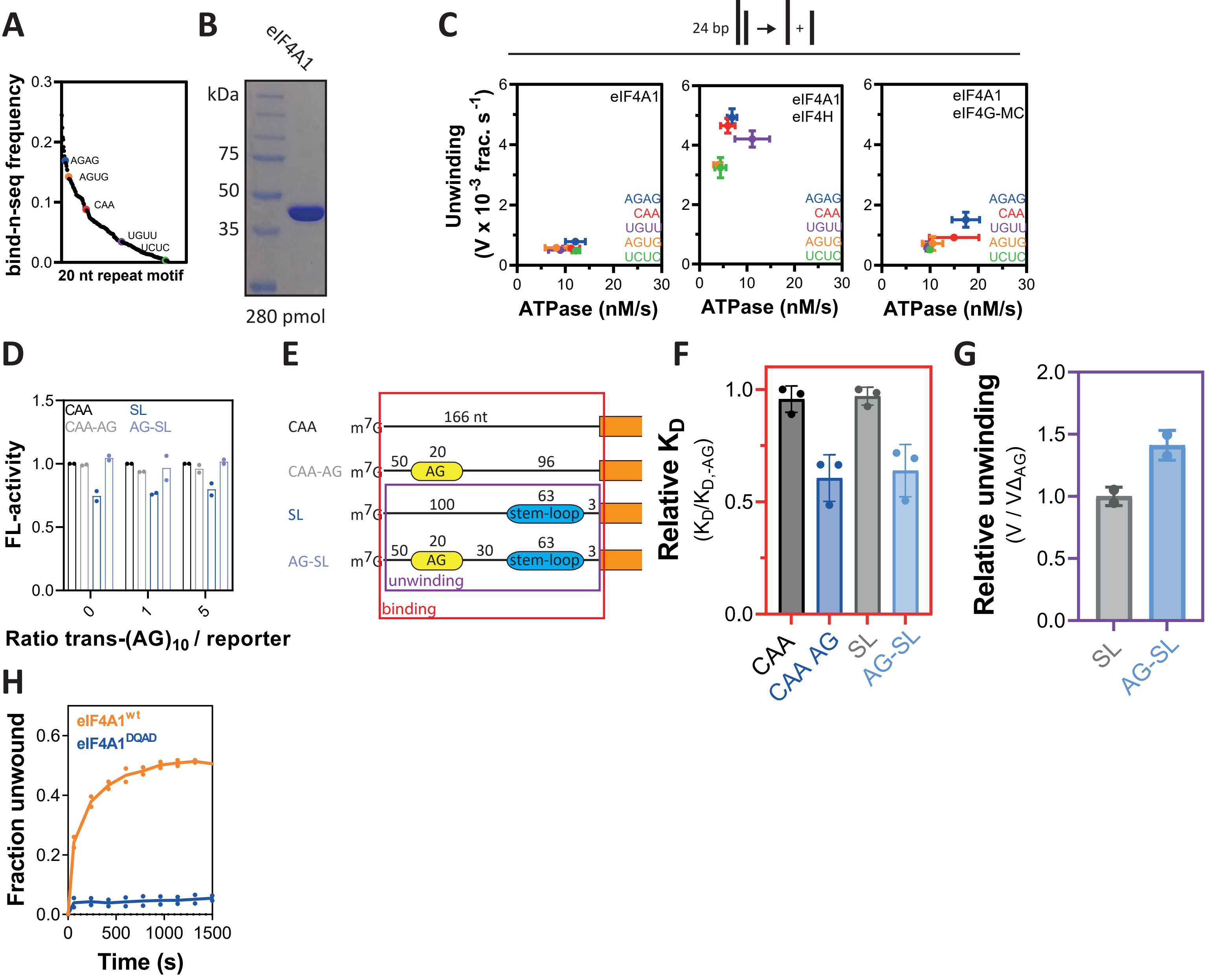


**Supplementary Figure 1. Unwinding by eIF4A1 is stimulated in an RNA sequence-dependent manner *in vitro*. A,** Relative binding affinity of eIF4A1 to 20 nt repeat ssRNA sequences measured as frequency of occurrence in a bind-n-seq study by Iwasaki et al. (Iwasaki et al., 2016) Figure has been created using data from Iwasaki et al. (Iwasaki et al., 2016) Highlighted motifs were used in this study. **B,** Coomassie-stained denaturing SDS-gel after electrophoresis of purified, recombinant eIF4A1. **C**, same as Fig. 1B, but including ATPase under the same conditions. Data are mean ± SEM from repeat experiments, n(ATPase) = 3 for each condition. **D,** Firefly luciferase activity after 60 min translation of reporters shown in Fig. 1C in rabbit reticulocyte lysate in the presence of equimolar or 5-fold excess of 20 nt AG-RNA. AG-RNA in *trans* to the reporter does not stimulate translation of SL-reporter nor affect translation the other reporters. Hence, effects described in Fig. 1D require the AG-repeat motif to be located in *cis* to the 5'UTR of the reporter mRNA. Data are mean from repeat experiments, n = 2. **E**, Indicated regions of the mRNA reporters in red and purple were generated and used as RNA substrates for RNA-binding experiments shown in Supplementary Fig. 1F and unwinding reactions shown in in Supplementary Fig. 1G, respectively. **F**, RNA-binding of eIF4A1 to RNA substrates described in Supplementary Fig. 1E in the presence of AMP-PNP. Data are mean ± sd from three repeat experiments. Binding of eIF4A1 to 5'UTR fragments was about 2-fold stronger if the 166 nt long 5'UTR fragment contained the 20 nt AG repeat sequence as shown in Supplementary Fig. 1E. **G**, unwinding of eIF4A1 using a 5' overhang 24bp substrate with 60 nt CAA-overhang with and without a 20nt AG-repeat 30 nt upstream the duplex region. The design of the substrate reflects the organisation of the 5'UTR fragment indicated in purple in Supplementary Fig. 1E. Data are mean ± sd from two repeat experiments Unwinding by eIF4A1 was enhanced if the RNA substrate contained the AG-repeat sequence, showing that the 20 nt AG-repeat in the 5'UTR recruits eIF4A1 and stimulates its unwinding activity. **H**, Progress curves of eIF4A1$^{wt}$ and eIF4A1$^{DQAD}$ on AG-overhang substrate confirming the unwinding-inactive state of eIF4A1$^{DQAD}$. Data are mean from a technical duplicate.

### Supplementary Figure 2

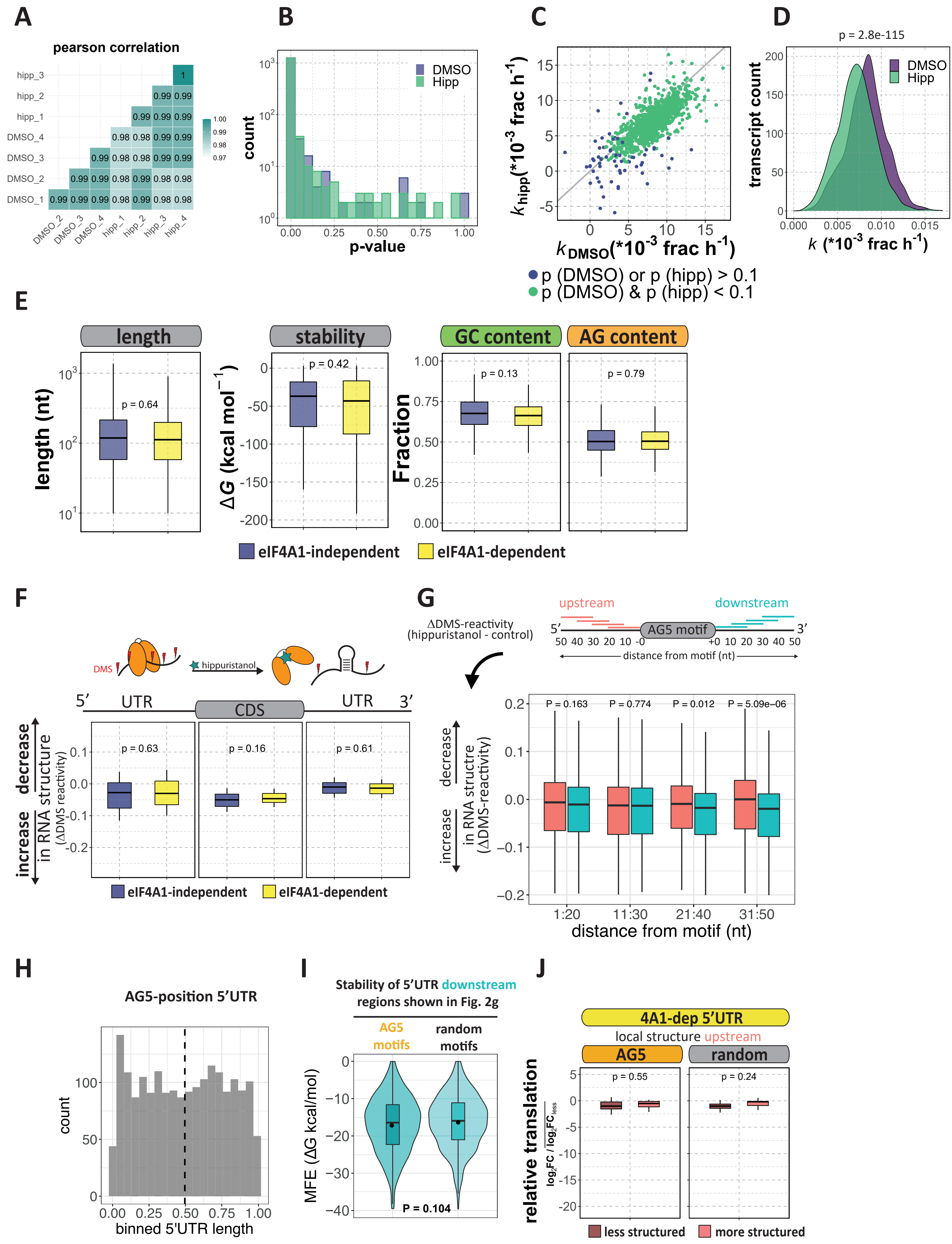

**Supplementary Figure 2. eIF4A1 shows RNA sequence-dependent unwinding and stimulation of translation in cells. A,** Pearson correlation coefficients between replicates of the TMT-pulsed SILAC from main figure 2, **B,** Histogram showing binned p-values (bins = 0.05) of the derived translation rates from the linear fit of the experimental TMT pulsed SILAC data from control DMSO and hippuristanol conditions. p-values were calculated from a F-test against k(slope) = 0. **C**, Scatter plot of translation rates under control (DMSO) or hippuristanol conditions grouped by p-value cut-off. Genes coloured in green (p-values <0.1, from fitting process supplementary Fig. 2B) were used for downstream analyses. n(green) = 1270, n(purple) = 67. **D**, Density plot of translation rates from DMSO control or hippuristanol conditions. In agreement with hippuristanol being a translational inhibitor, translation rates are globally downregulated upon eIF4A1 inhibition with hippuristanol. The p-value was calculated by a paired, two-sided Wilcoxon test. **E**, 5'UTR features of eIF4A1-dependent and –independent 5'UTRs mRNAs filtered for most abundant transcripts. 5'UTRs were folded and their mean free energy (Δ*G*) calculated using mfold server (Markham and Zuker, 2008; Zuker, 2003). Thermodynamic folding stabilities of the 5'UTRs of eIF4A1-dependent and –independent mRNAs are comparable. The p-values were calculated by unpaired, two-sided Wilcoxon tests. **F**, Box plots of the change in DMS-reactivity following eIF4A1 inhibition of the 5'UTR, CDS and 3'UTR of eIF4A1-dependent and –independent genes identified in the TMT-pSILAC. This shows that eIF4A1-dependent and -independent mRNAs experience a comparable change in RNA structure after eIF4A1-inhibition. This is in agreement with our previous report, that eIF4A1-inhibtion does not lead to global changes in RNA structure, but localized RNA structure (Waldron et al., 2019). The p-values were calculated by unpaired, two-sided Wilcoxon tests. **G**, upper panel, Schematic presentation of the conducted sliding window analysis. The DMS-reactivity of 20 nt sliding windows with 10 nt step size up- and downstream of 10 nt polypurine (AG5) or random motifs (control) have been compared pairwise. Lower panel, box plot of the average change in DMS-reactivity (ΔDMS reactivity) following eIF4A inhibition of all of 20 nt sliding windows down- and upstream of all 10 nt AG5 motifs in 5'UTRs. The Δ reactivity is calculated by subtracting the reactivity under control conditions from the reactivity under hippuristanol conditions, based on previous RNA Structure-seq2 data in MCF7 cells (Waldron et al., 2019). Therefore, a negative Δ reactivity indicates decreased reactivity following eIF4A inhibition, which is consistent with increased structure and *vice versa*. Changes in RNA structure around AG-motifs are most pronounced in a 20 nt window, 31 nt downstream of the AG motif which is also shown in Fig. 2F. The p-value was calculated by a paired, two-sided Wilcoxon test. **H**, Histogram depicting the binned position of all AG5 motifs in all 5'UTRs more than 100 nt in length from the most abundant transcript per gene in MCF7 cells based on Structure-seq2 data (Waldron et al., 2019). The dotted line depicts the median, which along with the bars indicates that R10 motifs are spread evenly throughout 5'UTRs. n = 1916 motifs from 1186 transcripts. **I**, Violin plots comparing the minimum free energy of the same 5'UTR windows shown in Fig. 2F. Dot is the mean; box plot shows the median. The p-value was calculated by a paired, two-sided Wilcoxon test. **J,** Box plots of the log2-fold change in translation rate following eIF4A1 inhibition of mRNAs containing 5'UTR AG5 motifs (or random control motifs) that show changes in local RNA structure 50 nt upstream the motif (same windows as Fig. 2F). Note, that Fig. 2G shows corresponding downstream RNA regions as defined in Fig 2F. The p-value was calculated by a paired, two-sided Wilcoxon test.

### Supplementary Figure 3

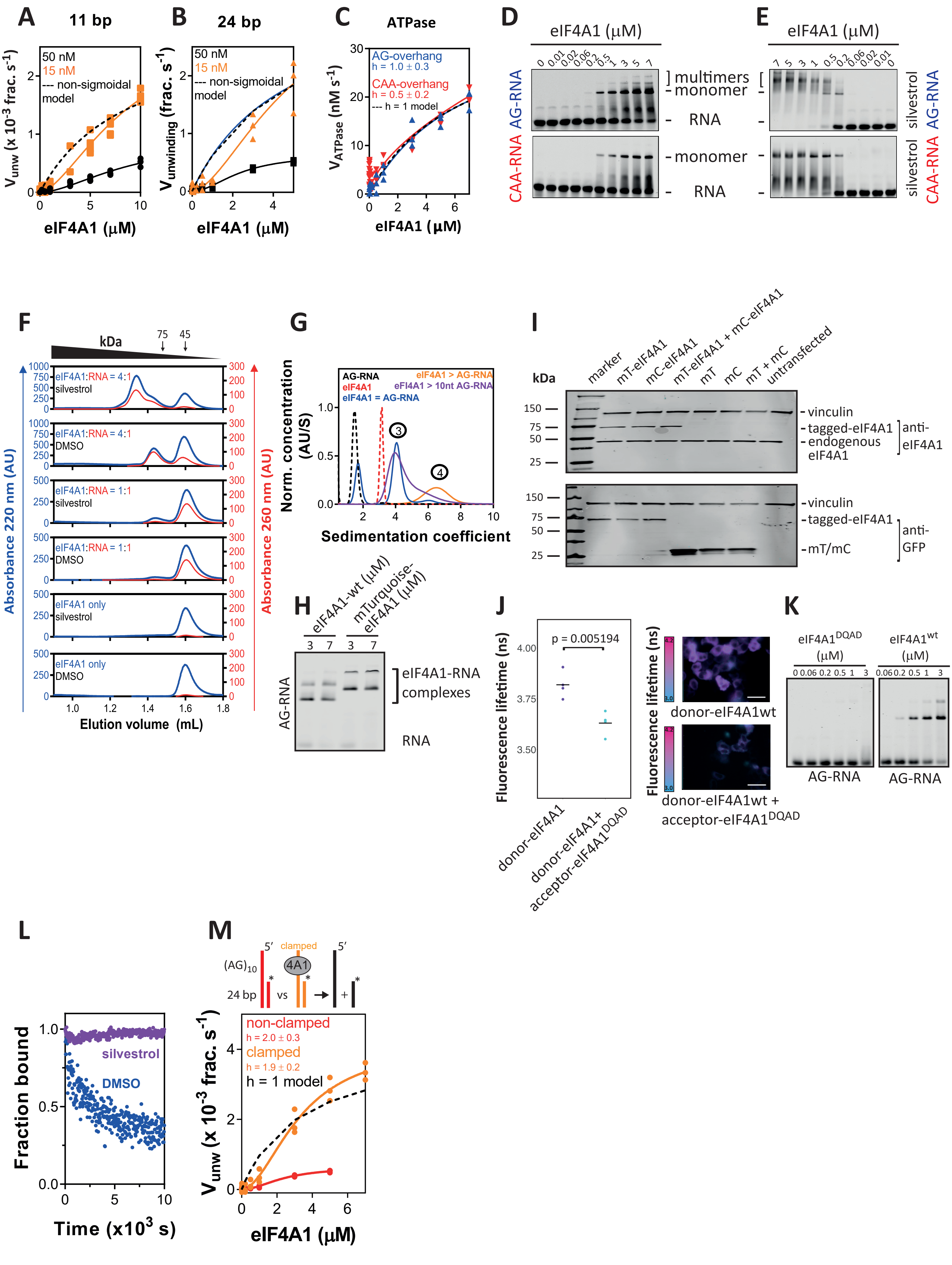

**Supplementary Figure 3. RNA sequence-specific unwinding by eIF4A1 is performed by a multimeric RNA-eIF4A1 complex. A-B,** Unwinding activity of eIF4A1 on a 20 nt AG-overhang substrate with 11 bp or 24 bp duplex region at indicated substrate concentrations. In all cases, a non-hyperbolic but sigmoidal shape of the curve is observed suggesting that independent of the duplex length unwinding by eIF4A1 involves more than one copy of the protein; data are mean (of technical duplicates) from three repeat experiments, n = 3. The dashed lines represent a hypothetical model with h = 1 in which eIF4A1 follows non-sigmoidal kinetics for a monomeric enzyme using the kinetic parameters from fitting the Hill-equation to the experimental data. **C,** ATPase activity, n(repeat experiments) = 3, of eIF4A1 on AG- and CAA-overhang substrate. The Hill-equation was fitted to the data. In contrast to the unwinding reactions shown in main Fig. 3A, the ATPase activity of eIF4A1 shows a non-cooperative behaviour. **D**, Representative electrophoretic mobility shift assays (EMSA) of 20 nt AG- or CAA-RNA at increasing eIF4A1 concentrations. n = 3 repeat experiments. **E,** Representative electrophoretic mobility shift assays of eIF4A1-binding to 25 nM AG- and CAA-RNA in the presence of silvestrol at increasing concentrations of eIF4A1. Multimeric eIF4A1-RNA complexes are detectable only with the AG-RNA; n = 3 repeat experiments. **F**, Analytical gel filtrations of eIF4A1-AG-RNA complexes at saturating protein (16 µM eIF4A1, 4 µM RNA) or RNA concentrations (4 µM eIF4A1, 12 µM RNA) in the presence or absence of silvestrol. Reference protein: Ovalbumin – 45 kDa at 1.6 mL, Conalbumin – 76 kDa at 1.5 mL, free RNA at 2.05 mL (not shown). **G,** Analytical ultracentrifugation of eIF4A1-AG-RNA complexes. eIF4A1 at 8 µM and RNA at 2 (eIF4A1>AG-RNA) or 8 µM (eIF4A1=AG-RNA). Data are summarised in Supplementary Table 2. This confirms results from the gel filtrations that eIF4A1-multimerisatoin is only induced upon RNA-binding. In addition, multimerisation is observed even on the short 10 nt AG-RNA suggesting that in multimeric eIF4A1 only one eIF4A1 molecule is in tight contact with the RNA as shown in a recent crystal structure of eIF4A1 in complex with the same 10 nt AG-RNA in the presence of rocaglamide A, a silvestrol derivate (Iwasaki et al., 2019). Modelling of the data (Supplementary Table 2) suggests a 3:1 stoichiometry of eIF4A1:AG-RNA in the multimeric complexes. **H,** EMSA comparing multimerisation of $eIF4A1^{wt}$ and mTurquoise-eIF4A1 upon binding to AG-RNA. The N-terminal fusion of mTurquoise to eIF4A1 does not affect its ability to form multimeric complexes. **I,** Western blots corresponding to Fig. 3D-E showing expression levels of overexpressed mTurquoise- (mT) and mCitrine-tagged (mC) eIF4A1 (anti-GFP blot) versus endogenous eIF4A1 (anti-eIF4A1 blot) expression. Vinculin is used as the loading control. uncropped blots are shown. **J**, FLIM-FRET images and quantification of co-transfection of mTurquoise-eIF4A1-wildtype and mCitrine-$eIF4A1^{DQAD}$; n = 4 independent repeat experiments. Total cells included all replicates (DQAD) = 129, (wt) = 171; p-value was calculated by an unpaired, two-tailed t-test. The scale is 50 µm. Cotransfection of tagged $eIF4A1^{wt}$ and $eIF4A1^{DQAD}$ yielded a similar change in fluorescence lifetime as with $eIF4A1^{wt}$ only (see Fig. 3E) indicating that RNA-binding capacity of at least one of the subunits in multimeric eIF4A1 is not essential for multimerisation. Hence a direct eIF4A1-eIF4A1 interaction induced by binding of one eIF4A1 to RNA is suggested. **K**, AG-RNA binding of $eIF4A1^{wt}$ and $eIF4A1^{DQAD}$ confirming the RNA-binding deficiency of $eIF4A1^{DQAD}$. **L,** ATPase-dependent RNA release of eIF4A1-AG-RNA complexes ± silvestrol. 4 µM protein were pre-incubated for 60 min with 1 µM AG-RNA in the presence or absence of 100 µM silvestrol together with ATP in the absence of magnesium. Then magnesium chloride was added to a final concentration of 2 mM and after 10 min incubation real-time fluorescence was recorded. Data are the mean of a technical duplicate. AG-RNA is not released in the presence of silvestrol while in the absence of silvestrol the RNA is released as a result of ATP hydrolysis. This shows that silvestrol clamps eIF4A1 to the AG-RNA in an ATP-independent manner as observed previously (Iwasaki et al., 2016). **M**, Unwinding activity of clamped and non-clamped eIF4A1 on 20 nt AG-overhang substrate. The Hill-equation was fitted to the data. Data are mean (of technical duplicates) from three repeat experiments. The back line models the experimental data assuming a non-cooperative (h = 1) mechanism. This shows that clamping eIF4A1 to the overhang stimulates unwinding when eIF4A1 is in excess over the RNA and that even under clamping conditions eIF4A1 unwinds RNA as a multimeric enzyme because h > 1.

### Supplementary Figure 4

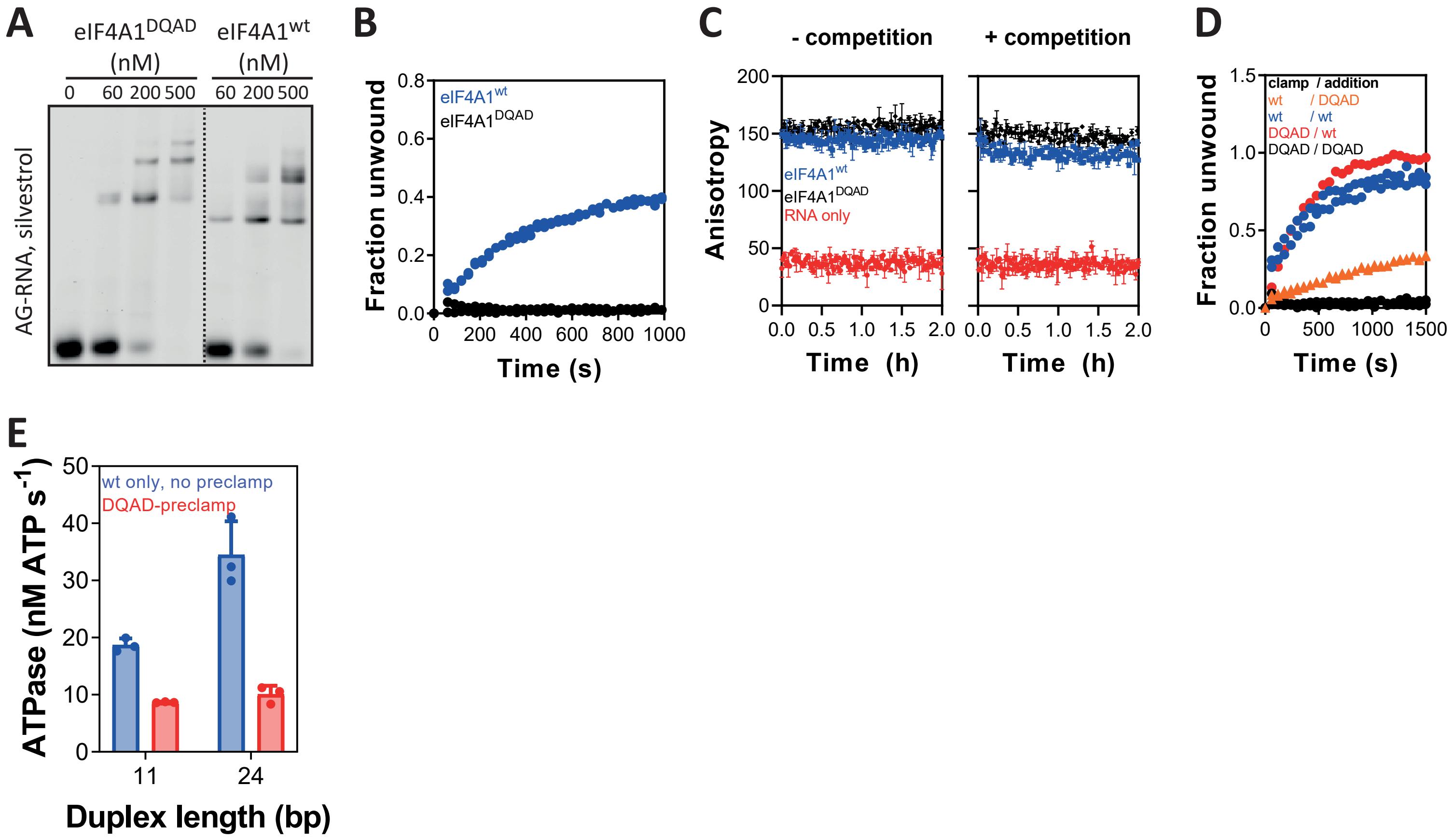


**Supplementary Figure 4. Division of functions in multimeric eIF4A1. A,** EMSA of AG-RNA binding and **B**, progress curve of unwinding of eIF4A1wt and eIF4A1$^{DQAD}$ both in the presence of silvestrol. A technical duplicate is shown. eIF4A1$^{DQAD}$ is catalytically inactive while it binds AG-RNA at similar affinity as eIF4A1wt in the presence of silvestrol. **C**, Representative progress curves (of three replicates) of AG-RNA release of eIF4A1$^{wt}$ and eIF4A1$^{DQAD}$ in the presence of ATP and silvestrol -/+ unlabelled competitor AG-RNA under conditions equivalent to the experiment shown in Figure 4B. The mean ± sd of a technical duplicate is shown. This shows that RNA-complexes of both wildtype and variant eIF4A1 display very similar stabilities under conditions equivalent to those in Figure 4B, arguing against eIF4A1$^{wt}$ replacing overhang-bound eIF4A1$^{DQAD}$. Thus, variant and wildtype protein cooperate to achieve unwinding as shown in Fig. 4B **D**, Progress curves of unwinding by indicated mixes of eIF4A1$^{wt}$ and eIF4A1$^{DQAD}$ at a total concentration of 5 µM following the reaction scheme shown in Fig 4B. A technical duplicate is shown. **E,** ATPase activities of unwinding competent multimeric eIF4A1 complexes comparing eIF4A1$^{wt}$ complexes with ones in which inactive eIF4A1$^{DQAD}$ was pre-clamped to the overhang region. Data are mean ± sd from three repeat experiments; n = 3. The result shows that the ATPase is stronger when eIF4A1$^{wt}$ is bound to the overhang. As the chimeric eIF4A1wt-eIF4A1$^{DQAD}$ complex also unwinds RNA efficiently (Fig. 4B), this suggests that the ATPase activity of actively unwinding eIF4A1 is lower than the eIF4A1-loading subunits that are bound to the overhang region.

### Supplementary Figure 5

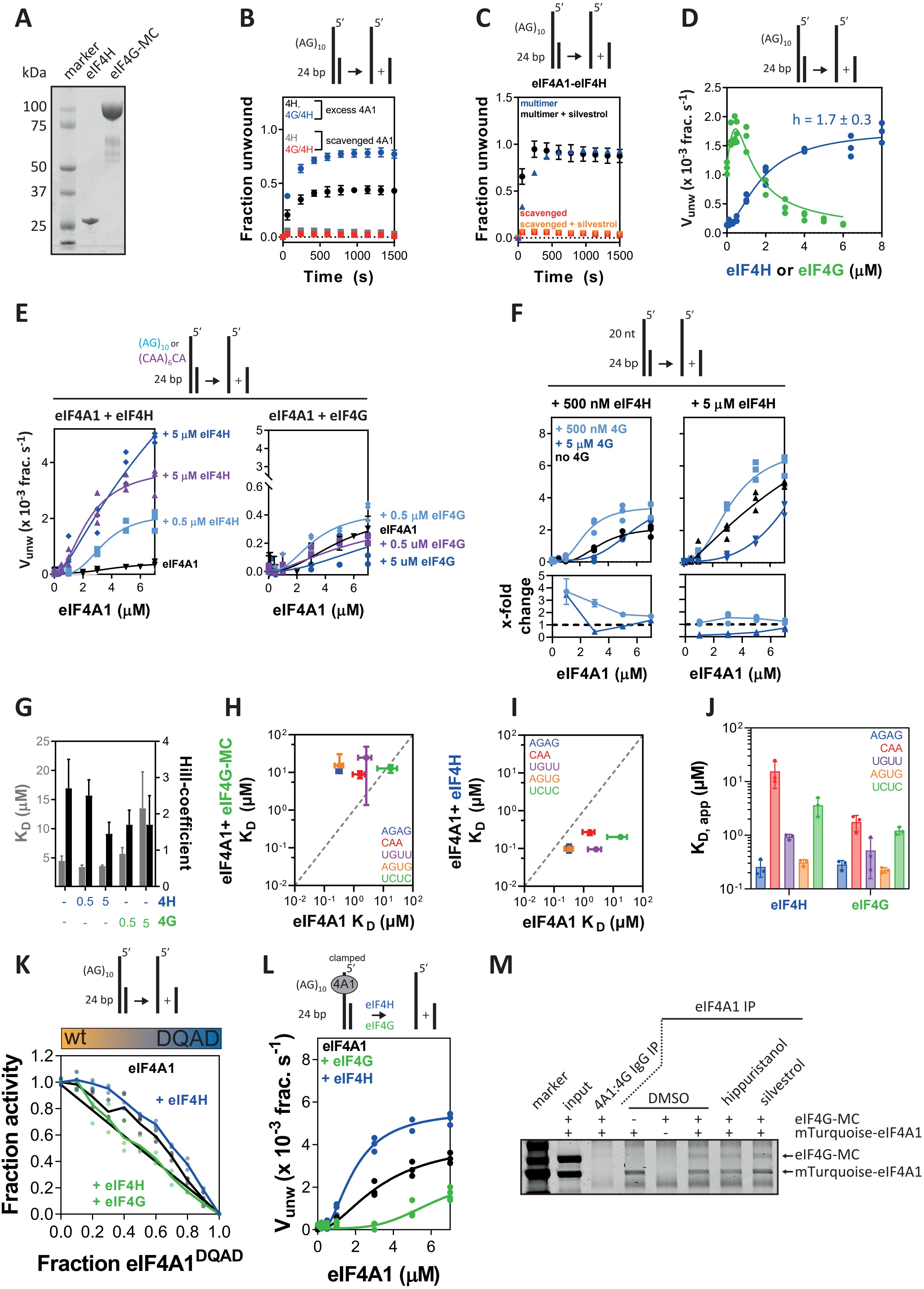

**Supplementary Figure 5. eIF4A1-cofactors function upon distinct eIF4A1 subunits within multimeric eIF4A1**. **A,** Coomassie-stained denaturing SDS-gel after electrophoresis of recombinant eIF4H and eIF4G-MC. **B,** Progress curves of 5 µM eIF4A1 on AG-overhang substrate in the presence of 500 nM eIF4H or 500 nM eIF4H and 500 nM eIF4G under excess or scavenged eIF4A1 conditions (See Fig. 3F). Data points are the mean of a technical duplicate ± sd. Unwinding by eIF4A1 is enhanced under excess eIF4A1 conditions even in the presence of the cofactors. **C,** Progress curves of 5 µM eIF4A1 on AG-overhang substrate in the presence of 5 µM eIF4H in the presence or absence of silvestrol. Data points are the mean of a technical duplicate ± sd. Regardless of clamping eIF4A1 to the substrate, unwinding is enhanced when excess eIF4A1 is present, suggesting that under 'scavenging' conditions RNA substrate and cofactor binding do not compete for eIF4A1. **D,** Unwinding activity of 2 µM eIF4A1 (blue) or 2 µM eIF4A1-eIF4H (1:1, green) at increasing concentrations of eIF4H (blue) or eIF4G (green), respectively; data are mean (of technical duplicates) from three repeat experiments; n = 3. The Hill-equation was fitted to the eIF4H-data and an equation describing substrate inhibition was fitted to the eIF4G-data (lines). **E,** Unwinding activity of eIF4A1 on AG- (blue) or CAA-overhang (purple) substrate in the presence of indicated concentrations of eIF4H (left panel) or eIF4G (right panel). Data are mean (of technical duplicates) from three repeat experiments; n=3. The Hill-equation was fitted to the data (lines). **F,** Unwinding activity of eIF4A1 on AG-overhang substrate in the presence of 500 nM (left panel) or 5 µM eIF4H (right panel) together with indicated concentrations of eIF4G. Data are mean (of technical duplicates) from three repeat experiments, n =3. The Hill-equation was fitted to the data (lines). The bottom panel show the change in unwinding activity as compared to in the absence of eIF4G. This shows that only when both cofactors are limiting synergistic stimulation of eIF4A1 by the cofactors is optimal. **G**, Hill-coefficient and functional binding affinities as derived from the fits to the experimental data shown in Supplementary Figure 5E. Bars show result and errors of the fit. Both cofactors lower the Hill-coefficient of unwinding by eIF4A1. Only eIF4G also lowers the apparent functional binding affinity indicating competition with eIF4A1 for substrate binding. **H**, RNA binding of eIF4A1 alone versus RNA-binding of eIF4A1 in the presence of eIF4G-MC or **I**, the presence of eIF4H; data are mean ± sd from three repeat experiments; n = 3. eIF4G reduces eIF4A1-RNA binding while eIF4H stimulates eIF4A1 RNA-binding activity. This confirms that eIF4G competes with eIF4A1 for RNA-binding. **J,** RNA-binding affinities of eIF4G and eIF4H to indicated RNAs. Data are mean ± SD from three repeat experiments, n =3. **K,** Inhibition of unwinding of eIF4A1$^{wt}$ by fractional mixes with eIF4A1$^{DQAD}$ alone (black) in the presence of 5 µM eIF4H (blue) or 2 µM eIF4H with 1 µM eIF4G (green) corresponding to the points of maximum stimulation as shown in Supplementary Fig. 5D. Activity data of each replicate is plotted relative to non-inhibited eIF4A1 (no added eIF4A1$^{DQAD}$); data are mean (of technical duplicates) from three repeat experiments, n = 3. The line indicates the behaviour of a monomeric or multimeric enzyme without subunit-cooperativity. In the presence of eIF4H the cooperation between eIF4A1 subunits is maintained, while in the presence of eIF4G eIF4A1-subunit cooperativity is lost. As unwinding is under all conditions still performed in a cooperative manner (Supplementary Fig 5E-F), this suggests that eIF4G is coordinating the activity of the eIF4A1 unwinding subunits. This might be achieved by eIF4G replacing the loading subunit of multimeric eIF4A1. **L,** Unwinding activity of clamped eIF4A1 on AG-overhang substrate alone (black) and in the presence of 5 µM eIF4H (blue) or eIF4G (green). Data are mean (of technical duplicates) from three repeat experiments, n = 3. The Hill-equation was fitted to the data (lines). This shows that eIF4H stimulates clamped eIF4A1 while eIF4G is inhibitory to the activity of clamped eIF4A1 suggesting that eIF4H functions upon the unwinding-subunits of eIF4A1 while eIF4G operates upon or replaces the loading-subunits of multimeric eIF4A1. **M,** Coomassie-stained SDS-gel of coimmunoprecipitated recombinant mTurquoise-eIF4A1 and eIF4G-MC using an anti-eIF4A1 antibody in the presence of indicated eIF4A1 inhibitors. None of the eIF4A1-inhibitors disrupt the eIF4A1-eIF4G interaction.

### Supplementary Figure 6

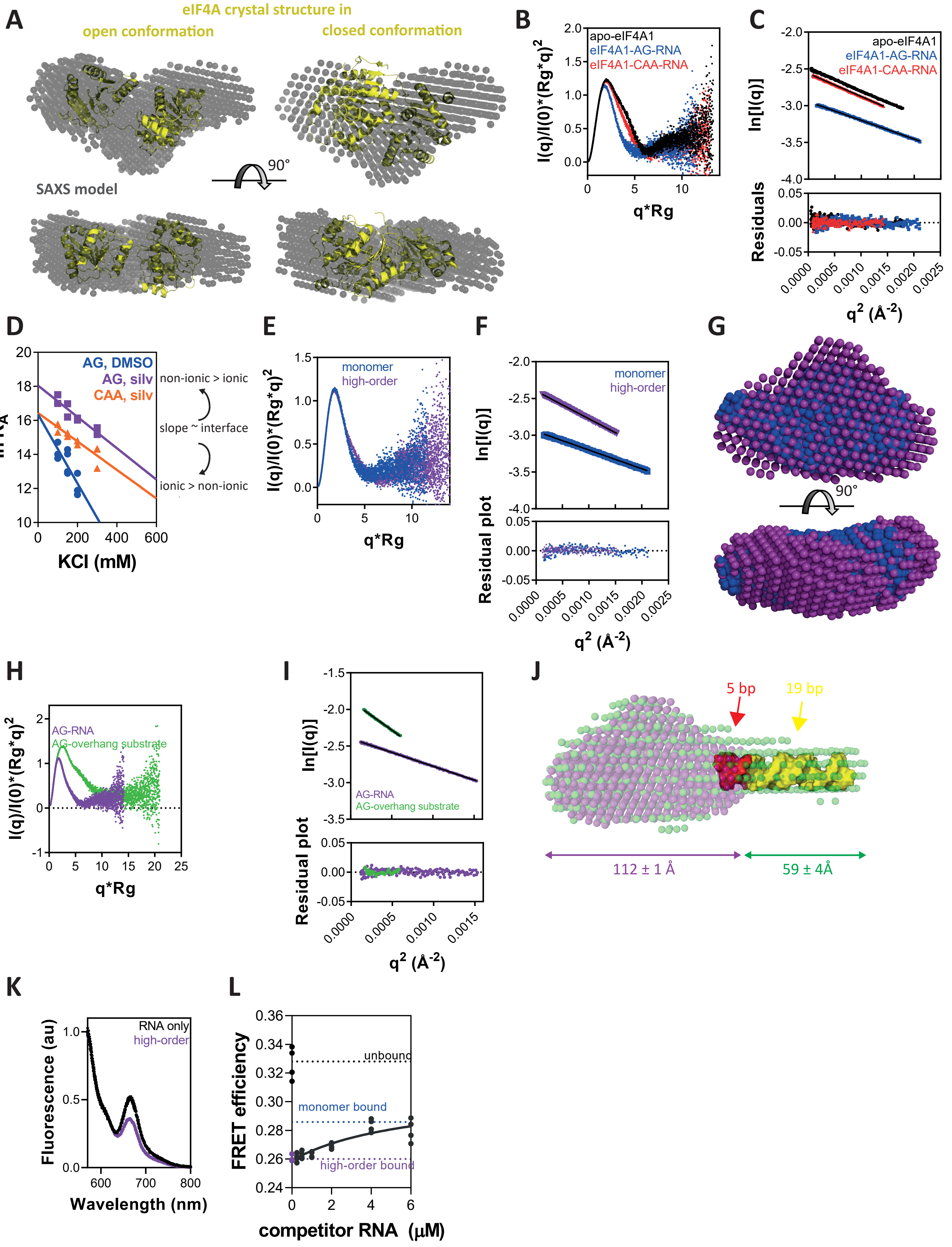

**Supplementary Figure 6. The architecture of eIF4A1-loading complexes. A,** Superposition of the envelope model of apo-eIF4A1 (grey) derived from our SAXS data with the crystal-structure of *S. cerevisiae* eIF4A (yellow) representative of an half-open conformation of eIF4A (PDBid 2vso (Schutz et al., 2008), $\chi^2 = 10$), and to the crystal structure of eIF4A1 in complex with a 10nt AG-RNA which is representative of the closed conformation (PDBid 5zc9 (Iwasaki et al., 2019), $\chi^2 = 163$). This suggests that the derived SAXS-model for solution apo-eIF4A1 is more representative of an open and more dynamic conformation than a closed and compact one. **B,** Dimensionless Kratky and **C** Guinier plots of SAXS data corresponding to apo-eIF4A1 and monomeric eIF4A1 bound to AG or CAA-RNA as shown in Fig. 5A. The Kratky plot demonstrates that, while all proteins are in a folded and globular state (bell shape), the eIF4A1-AG-RNA complex is more compact than the CAA-RNA complex and the apo-eIF4A1 (broader curve of CAA-eIF4A1 and apo-eIF4A1 *vs* AG-eIF4A1). The slope of the linear fit in the Guinier plot is used to calculate the radius of gyration. The good quality of the linear fit to the data in the Guinier plot shows negligible aggregation in the samples. **D,** Linear free energy relationships (LFER) between the association constant ($K_A = 1/K_D$) of eIF4A1 to AG- and CAA-RNA and increasing potassium chloride concentrations in the presence or absence of silvestrol. Data are mean ± sd from at least two repeat experiments ($n \geq 2$). The slopes of the linear fits are measures for the relative contribution of ionic and non-ionic interactions in the binding process (Schmidt et al., 2016). Our results show that the addition of silvestrol reduced the dependency of eIF4A1 binding to AG-RNA on the salt concentration. Hence, in the presence of silvestrol, there are additional non-ionic interactions contributing to the RNA-protein interface than in the absence of silvestrol. This is in agreement with the recent crystal structure of eIF4A1 and a 10 nt AG-repeat RNA in the presence of the silvestrol-derivate rocaglamide (Iwasaki et al., 2019) that reveals hydrophobic base stacking interactions between the RNA and the eIF4A1-rocA complex bridged by rocA. The LFER slopes of eIF4A1 binding to AG and CAA RNA are similar in the presence of silvestrol suggesting a comparable contribution of ionic interactions between these two eIF4A1-RNA complexes. As the $K_A$ are different, this accordingly suggests that there is a higher fraction of non-ionic interactions present in the eIF4A1-AG-RNA interface than in the eIF4A1-CA-RNA interface. Note, that we could not measure a LFER for CA-RNA-eIF4A1 in the absence of silvestrol accurately as affinities were very weak at increased salt concentrations. **E,** Dimensionless Kratky and **F** Guinier of SAXS data corresponding to monomeric and multimeric eIF4A1 as shown in Fig 5B. The good quality of the linear fit to the data in the Guinier plot shows negligible aggregation in the samples. **G**, Superposition of the SAXS models corresponding to the monomeric (blue) and multimeric (purple) eIF4A1-AG-RNA complexes. **H,** Dimensionless Kratky and **I** Guinier of SAXS data corresponding to multimeric eIF4A1 bound to AG-ssRNA or AG-overhang substrate as shown in Fig 5D. The Kratky plot of the eIF4A1-overhang substrate complex shows that is has a more extended shape than the eIF4A1-AG-RNA complex and indicates that it has two diffracting domains as suggested by the shoulder in the curve. The good quality of the linear fit to the data in the Guinier plot shows negligible aggregation in the samples. **J**, A 24 bp dsRNA (yellow, extracted from PDBid 2L3J) was manually fitted into the experimental SAXS envelope of multimeric eIF4A1 bound to the overhang of a 24 bps duplex substrate using PyMOL2. This suggests that multimeric eIF4A1 not only binds the single stranded overhang region of the RNA substrate but also, in addition, is directly located at the ssRNA-duplex fork and may cover ~ 5 bps of the duplex region which depicts the hypothetical priming mechanism of unwinding. **K,** Representative (of four repeat experiments) fluorescence-emission spectra (excitation at 520 nm) of FRET-labelled AG-overhang substrate in its unbound/free state (black) or bound by multimeric eIF4A1 (purple). **L,** Relative FRET efficiency of AG-overhang substrate when bound to eIF4A1 derived from fluorescence emission spectra (example shown in Supplementary Figure 6K); data from four repeat experiments; n = 4. Binding of multimeric eIF4A1 to AG-overhang region induces a specific conformational change in the ssRNA-region of the substrate that is different from eIF4A1-monomer
